# Systematic morphological profiling of human gene and allele function reveals Hippo-NF-κB pathway connectivity

**DOI:** 10.1101/092403

**Authors:** Mohammad H Rohban, Shantanu Singh, Xiaoyun Wu, Julia B Berthet, Mark-Anthony Bray, Yashaswi Shrestha, Xaralabos Varelas, Jesse S Boehm, Anne E Carpenter

**Affiliations:** Broad Institute of MIT and Harvard, Cambridge, Massachusetts, USA; Department of Biochemistry, Boston University School of Medicine, Boston MA USA; Present address: Novartis Institutes for BioMedical Research, Cambridge MA USA

## Abstract

We hypothesized that human genes and disease-associated alleles might be systematically functionally annotated using morphological profiling of cDNA constructs, via a microscopy-based Cell Painting assay. Indeed, 50% of the 220 tested genes yielded detectable morphological profiles, which grouped into biologically meaningful gene clusters consistent with known functional annotation (e.g., the RAS-RAF-MEK-ERK cascade). We used novel subpopulation-based visualization methods to interpret the morphological changes for specific clusters. This unbiased morphologic map of gene function revealed TRAF2/C-REL negative regulation of YAP 1/WWTR1-responsive pathways. We confirmed this discovery of functional connectivity between the NF-**κ**B pathway and Hippo pathway effectors at the transcriptional level, thereby expanding knowledge of these two signaling pathways that critically regulate tumor initiation and progression. We make the images and raw data publicly available, providing an initial morphological map of major biological pathways for future study.

## Introduction

The dramatic increase in human genome sequence data has created a significant bottleneck. The number of genes and variants known to be associated with most human diseases has increased dramatically (Amberger et al. 2015). Unfortunately, the next step - understanding the function of each gene and the mechanism of each allele in the disease - typically remains non-systematic and labor-intensive. Most commonly, researchers painstakingly design, develop, and apply a disease-specific or biological process-specific assay.

Over 30% of genes in the human genome are of unknown function (Leonetti et al. 2016) and even annotated genes have additional functions yet to be uncovered. Furthermore, even when a gene’s normal functions are known, methods are lacking to predict the functional impact of the millions of genetic variants found in patients. These gaps must be filled in order to convert the promise of human genome sequence data into clinical treatments.

Therefore, there is a widespread need for systematic approaches to functionally annotate genes and variants therein, regardless of the biological process or disease of interest. One general approach depends on guilt-by-association, linking unannotated genes to annotated ones based on properties such as protein-protein interaction data, sequence similarity, or, most convincingly, functional similarity (Shehu, Barbará, and Molloy 2016). In the latter category are profiling techniques, where dozens to hundreds of measurements are made for each gene perturbation and the resulting profile is compared against profiles for annotated genes. Various data sources can be used for profiling; gene expression is one that can be performed in relatively high-throughput and it has been proven useful in predicting gene function (Lamb et al. 2006). In fact, high-throughput mRNA profiles were recently used to cluster alleles found in lung adenocarcinoma based on their functional impact, a precursor to therapeutic strategy for variants of previously unknown significance (Berger et al. 2016).

Images are a less mature data source for profiling but show tremendous promise. Morphological profiling data is complementary to transcriptional profiling data (Wawer et al. 2014) and is less expensive. Morphological profiling has succeeded across several applications, including grouping small-molecule perturbations based on their mechanism of action (Caicedo, Singh, and Carpenter 2016; Bougen-Zhukov et al. 2016), and grouping genes based on morphological profiles derived from cells perturbed by RNA interference (RNAi) (Mukherji et al. 2006; Boutros and Ahringer 2008; Fuchs et al. 2010; Pau et al. 2013). One limitation of RNAi for morphological profiling is that the number of measurements must be limited or else the resulting profiles are dominated by off-target effects, especially seed effects (Singh et al. 2015). Some computational solutions have shown some promise in overcoming this problem for gene expression profiling (Schmich et al. 2015), but their utility is unproven for image-based profiling, and regardless RNAi does not permit analysis of gene variants, only knockdown. Modification of genes via CRISPR will require new libraries of reagents and is as yet untested in morphological profiling.

In the proof-of-concept work presented here, we tested morphological profiling using overexpression in human cells as a general approach to annotate gene and allele function. We profiled a reference series of well-known genes, and a small number of variants thereof, by Cell Painting. In particular, we wondered whether the information content of this strategy would outweigh potential limitations (e.g., due to cellular context or expression level). We found that the approach successfully clustered genes and alleles based on functional similarity, revealed specific morphological changes even when present in only a subpopulation of heterogeneous cells, and uncovered novel functional connections between important biological pathways.

## Results

### Morphological profiles from Cell Painting of expression constructs are sensitive and reproducible

To profile each exogenously expressed gene (or allele therein), we used our previously developed image-based profiling assay, called Cell Painting (Gustafsdottir et al. 2013; Bray et al. 2016). This microscopy-based assay consists of six stains imaged in five channels and revealing eight cellular components: DNA, mitochondria, endoplasmic reticulum, Golgi, cytoplasmic RNA, nucleoli, actin, and plasma membrane (Fig. 1A). In five replicates in 384-well plate format, we infected U-2 OS cells (human bone osteosarcoma cells), chosen for their flat morphology and previous validation in the assay, with an arrayed “reference” expression library of 323 open reading frame (ORF) constructs of partially characterized functions (Supp. Table 1), a subset of which have been previously described (E. Kim et al. 2016). Of these, we prioritized analysis of the 220 constructs that were most closely representative of the annotated full length transcripts (see Methods). Morphological profiles were extracted using CellProfiler for image processing, yielding 1,384 morphological features per cell, and Python/R scripts for data processing, including feature selection and dimensionality reduction (Fig. 1B, and see Methods). This computational pipeline yielded a 158-dimensional profile for each of 5 replicates for each gene or allele tested.

**Figure 1:**
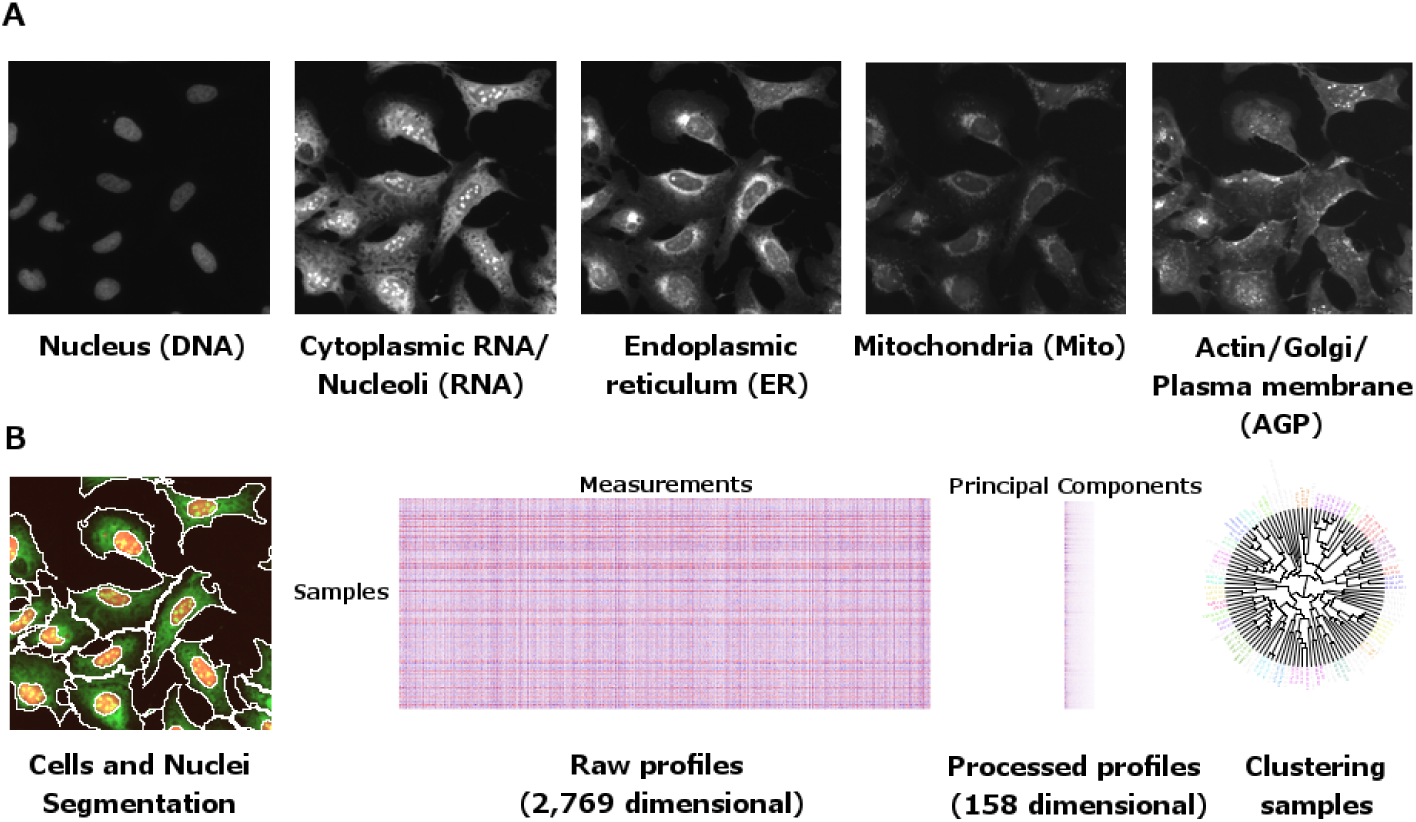
**Morphological profiling by Cell Painting**. (A) Example Cell Painting images from each of the five channels for a negative control sample (no gene introduced). (B) From left to right: Cell and nucleus outlines found by segmentation in CellProfiler; raw profiles (2,769 dimensional) containing median and median absolute deviation of each of 1,384 measurements over all the cells in a sample, plus cell count; processed profiles which are made less redundant by feature selection and Principal Component Analysis; dendrogram constructed based on the processed profiles (see Fig. 3). Replicates are merged to produce a profile for each gene which is then compared against others in the experiment to look for similarities and differences.

Not all genes are likely to impact cellular morphology given the limitation of our experiment-using a single cell line at a single time point under a single set of conditions and stained with six fluorescent labels. We therefore first asked what fraction of these ORFs impacted morphology. Surprisingly, we found that 50% (110/220) of these ORF constructs induced reproducible morphological profiles distinct from negative control profiles (Fig. 2A, and see Methods). Next, we ruled out the possibility that position artifacts may have artificially inflated this result by taking an alternative pessimistic null distribution which takes well position into account (Supp. Fig. 1). Therefore, we conclude that a single “generic” morphological profiling assay can detect signal from a substantial proportion of human genes. We next turned to testing whether those signals are biologically meaningful and can lead to novel, unbiased discoveries about gene function.

**Figure 2:**
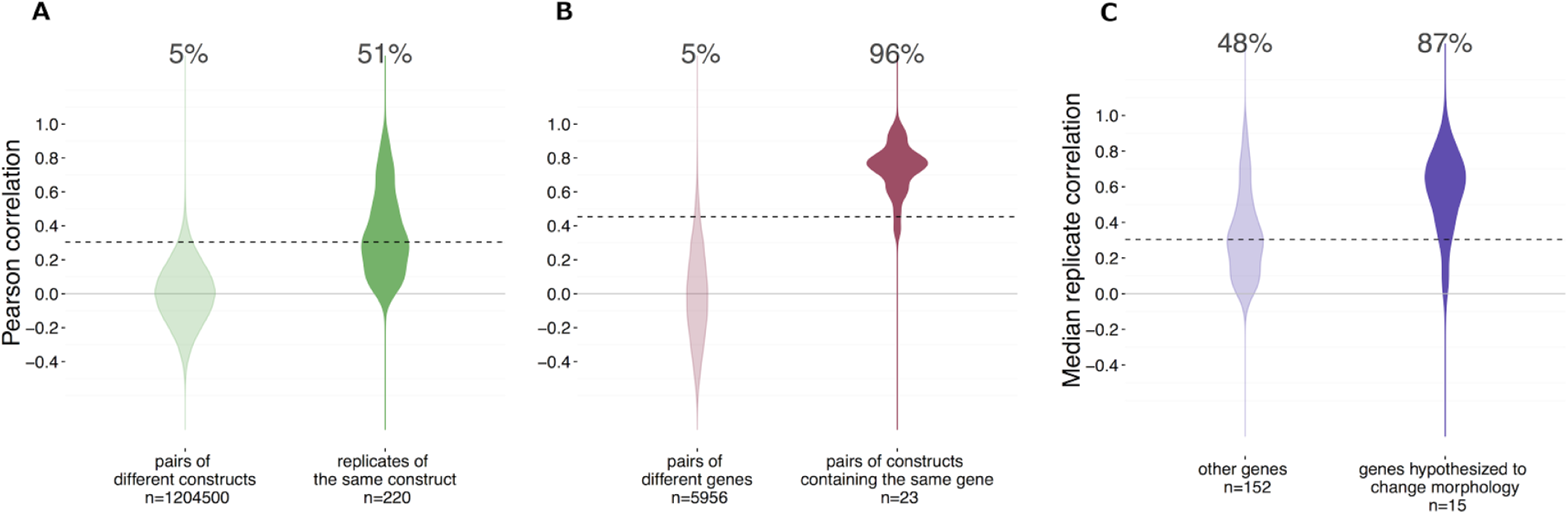
**(A) 50% of the gene overexpression constructs produced a detectable phenotype by image-based profiling**. Constructs yielding a reproducible phenotype ought to have a median correlation among replicates that is higher than the 95th percentile of correlations seen for pairs of different constructs; this is true for 51% (112 out of 220) of the constructs (as shown). Additionally, we removed two constructs that passed that filter but whose profiles were highly similar to negative control profiles (not shown), leaving 110 constructs (50%) for further analysis. **(B) Of wild-type ORF pairs that both yielded a distinguishable phenotype, 96% showed significant correlation to each other**. Correlations between the 23 pairs of constructs that are clones of the same gene (although with potential sequence variation or possibly different isoforms) were almost always much higher than correlations between pairs of constructs related to different genes. (C) **Genes in pathways thought to regulate morphology were more likely to yield detectable phenotypes** vs. the remainder of genes in the experiment (87% vs. 48%).

### Morphological profiling is robust, showing expected relationships

Given that technical replicates produce similar morphological profiles, we next evaluated whether similarities between profiles induced by different constructs are meaningful. We began with the simplest case: for a subset of genes in the experiment, a “wild-type” sequence (see Methods for important definitions) was captured in more than one ORF construct (23 pairs). These pairs either correspond to different physical cloning events and preparations but with highly similar full-length sequence (as defined in Methods; category a: 9 pairs), or a substantive difference in their nucleotide sequence, for example, isoforms (category b: 14 pairs). We found that, as expected, the phenotypes of over-expressed wild-type ORFs of the same gene were more similar to each other, on average, than to randomly selected genes. Of the 23 pairs for which both wild-type ORFs yielded a phenotype distinguishable from negative controls, 22 (~96%) of the pairs’ profiles were correlated more than expected by chance (Fig. 2B, the one pair not meeting that threshold was in category b), confirming that different constructs with biological similarity indeed produce similar morphological profiles.

This result also confirms that the sequence differences seen in separately cloned wild-type constructs do not generally have a major functional impact, but we caution that any individual construct of interest may have an impactful mutation; thus the raw sequence data should be examined and testing alternate constructs for a gene may be recommended. We also note that the 23 pairs analyzed here are located in different well locations on each plate; this result therefore also rules out widespread artifacts, such as plate position effects or metadata errors.

We suspected that the small number of engineered constitutively activating alleles for certain genes would, on average, yield a stronger phenotype than their wild-type counterparts. We indeed found that correlations between replicates of the constitutively activating allele were typically higher than correlations between replicates of the wild-type version of a gene (Supp. Table 2; p-value = 0.012, one-sided paired t-test).

We hypothesized that genes in pathways known to affect cellular morphology (RAC1, KRAS, CDC42, RHOA, PAK1, and Hippo) would be more likely to yield a morphological phenotype distinguishable from negative controls than other genes in the analysis. Indeed, we found this to be true (Fisher’s test p-value = 3.7 × 10^-3^) (Fig. 2C). Reassured by this validation, we were curious which pathways would be most and least likely to yield detectable morphological phenotypes, recognizing that “pathways” are neither separate nor well-defined entities. We found genes manually annotated as being in the Hippo, Hedgehog, cytoskeletal reorganization, and Mitogen-activated protein kinases (MAPK) pathways were more likely to result in a phenotype, whereas genes annotated as belonging to the JAK/STAT, hypoxia, and BMP pathways were among the least likely to yield a phenotype under the conditions tested (Supp. Fig. 2). Nevertheless, the majority of pathways could be interrogated by morphological profiling.

### Morphological signature similarity captures known gene-gene relationships

Given the caveats and limitations of overexpressing genes (see Discussion), we next tested whether image-based profiling of expression constructs could capture relationships among genes known to be functionally related. Because a reliable and complete map of all gene-gene connections is not available, we evaluated the accuracy of our results via two approaches.

First, we compared our data to protein-protein interaction data from BioGRID (Stark et al. 2006). This is imperfect ground truth for judging our predictions because two proteins might physically interact without producing the same morphological phenotype when overexpressed, and genes in the same pathway might regulate the same phenotype without any physical interaction. Nevertheless, we expect that the corresponding proteins of gene pairs with highest profile similarity are more likely than average to physically interact. Indeed, looking at wild-type versions of genes showing a detectable phenotype (the 73 genes represented in the 110 constructs), the ratio of verified gene connections among the top 5% correlated gene pairs (9%, 13 verified out of 143 possible combinations) is significantly higher than that of other gene pairs (5%, 128 verified out of 2,485 possible; Fisher’s test p-value = 0.04; Supp. Table 3).

Second, we manually annotated each gene for the pathway with which it is associated. This approach is based on expert opinion and thus imperfect knowledge of all genes’ function; furthermore many pathways interrelate, and genes in the same pathway are not expected to have identical phenotypes given that their functions are rarely identical (most notably, overexpression of some may activate while others suppress a biological pathway or process). Nonetheless, we expect pairs of genes whose morphological profiles correlate highly to be more likely than average to be annotated in the same pathway vs. different pathways. Using the same 73 genes as in the previous analysis, the ratio of gene connections with the same-pathway annotation in the top 5% most-correlated gene pairs was 20% (29 pairs out of 143), significantly higher than the ratio for the remaining pairs (6%, 139 pairs out of 2,485; Fisher’s test p-value = 7.53 x 10^-9^; Supp. Table 4).

### An initial morphological map of gene function

Having quantitatively established that morphological profiling is sensitive, robust, and captures known gene-gene relationships, we explored these relationships in a correlation matrix (Fig. 3 bottom left and Supp. Fig. 3). The overall structure, with multiple groupings along the diagonal, is consistent with the fact that the 110 constructs (73 unique genes) that showed a phenotype had been annotated as representing 19 different pathways. That is, we did not see large, homogeneous clusters, as would be expected if morphological profiling was sensitive to perturbation but not highly specific. This rules out uniform toxicity induced by a large number of genes, for example. Neither did we see only signal along the diagonal, which would have indicated no strong similarity between any gene pairs.

We next created a dendrogram (Fig. 3) and defined 25 clusters (see Methods and Supp. Fig. 4) to explore the similarities among genes. Pairs of wild-type ORFs almost always clustered adjacently, consistent with our quantitative analysis described above (Fig. 2B). We found that the majority of clusters (19 out of the 22 clusters containing more than one gene) were enriched for one or more Gene Ontology terms (Supp. Table 5), indicating shared biological functions within each cluster.

Using this dendrogram, we began by interrogating three clusters that conformed well to prior biological knowledge. First, we analyzed Cluster 20, containing the two canonical Hippo pathway members YAP1 and WWTR1 (more detail in Cluster 20A and 20B PDFs, and in a later section of the text). Both are known to encode core transcriptional effectors of the Hippo pathway (Johnson and Halder 2014), and a negative regulator of these proteins, STK3 (also known as MST2), is the strongest anti-correlating gene for the cluster (Cluster 20A PDF, panel c1).

Second, we noted Cluster 21 is comprised of the two phosphatidylinositol 3-kinase signaling/Akt (PI3K) regulating genes, PIK3R1 and PTEN, both frequently mutated across 12 cancer types in The Cancer Genome Atlas (TCGA) (Kandoth et al. 2013). These results are consistent with previous observations that certain isoforms of PIK3R1 reduce levels of activated Akt, a dominant negative effect (Abell et al. 2005) AKT3 itself is in the top five strongest anti-correlating genes for Cluster 21 (Cluster 21A PDF, panel c1).

**Figure 3:**
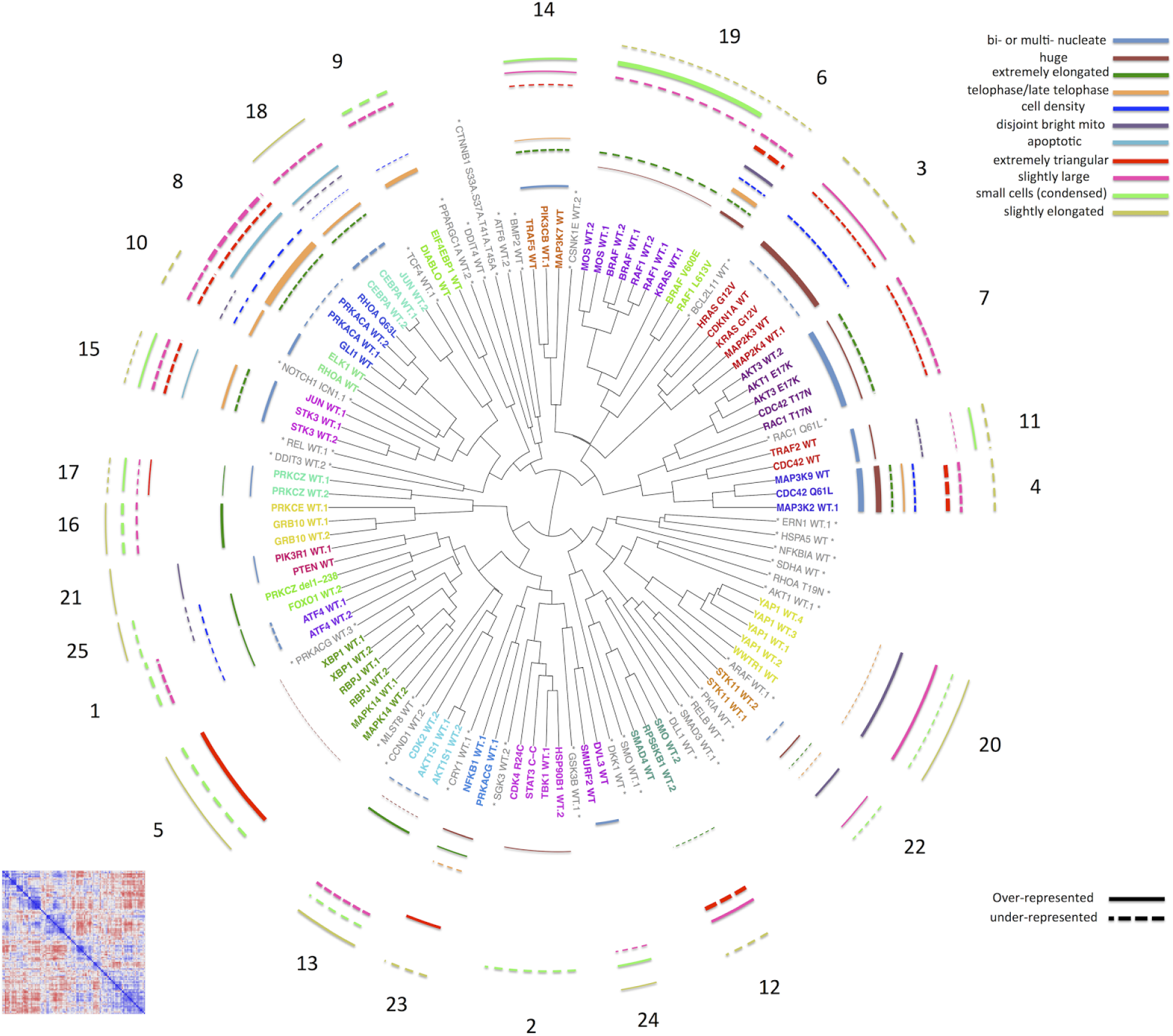
**Morphological relationships among overexpressed genes/alleles, determined by Cell Painting**. Correlations between pairs of genes/alleles were calculated and displayed in a correlation matrix (bottom left inset, full resolution is available as Supp. Fig. 3). Only the 110 genes/alleles with a detectable morphological phenotype were included. The rows and columns are ordered based on a hierarchical clustering algorithm such that each blue submatrix on the diagonal shows a cluster of genes resulting in similar phenotypes. The correlations were then used to create a dendrogram (main panel) where the radius of the subtree containing a cluster shows the strength of correlation. The 25 clusters containing at least two constructs are printed on the dendrogram in arbitrary colored fonts, while gene names colored gray and marked by asterisks are those that do not correlate as strongly with their nearest neighbors (i.e., they are singletons or fall below the threshold used to cut the dendrogram for clustering). Each colored arc corresponds to a cell subpopulation as noted in the legend. Line thickness indicates the strength of enrichment of the subpopulation in the cluster samples compared to the negative control. Solid vs. dashed lines indicate the over- vs. under-representation of the corresponding subpopulation in a cluster, respectively. Note that the number next to each cluster in the dendrogram is referenced in the main text and corresponds to the numbered supplemental data file for each cluster.

Third, we examined three Clusters (19, 6 and 3) that included many MAPK-related genes. Cluster 19 is the largest example of a tight cluster of genes already known to be associated; it includes four activators in the RAS-RAF-MEK-ERK cascade: KRAS, RAF1 (CRAF), BRAF, and MOS. Notably, two constitutively active alleles of these genes, BRAF^V600E^ (H. Davies et al. 2002) and RAF1^L613V^ (Wu et al. 2011), form a separate cluster (Cluster 6) adjacent to their wild-type counterparts. Furthermore, the constitutively active RAS alleles HRAS^G12V^ and KRAS^G12V^ (McCOY, Bargmann, and Weinberg 1984) are in the next-closest cluster (Cluster 3), which also contains MAP2K4 and MAP2K3 (known to be activated by Ras (Shin et al. 2005)), as well as CDKN1A (Jalili et al. 2012). By contrast, MAPKs that are known to be unrelated to the RAS-RAF-MEK-ERK cascade, such as MAPK14 in Cluster 5, are far away in the dendrogram.

Overall, these results support the notion that connections between genes can be efficiently discovered using our approach.

### Visualization approaches to assist interpretation of morphological signatures

We hypothesized that the specific morphologic features that segregated each of the clusters would provide insight into gene function. Examining images (Cluster 19A PDF, panel 3) or rank-ordered lists of features that distinguish individual profiles or clusters (Supp. Table 6) is tedious and lacks sensitivity for all but the most obvious of phenotypes, confirming that quantitative morphological profiling is more sensitive than the human visual system.

We therefore devised several strategies to enhance biological interpretability from these experiments and applied these in combination. First, we grouped features into meta-features based on their type of measurement, i.e., shape, texture, intensity, etc., and the cell constituents to which they are related, to create a Feature Grid (Fig. 4A). Second, we performed unsupervised grouping of features by mapping the top 20 most-distinguishing features for each cluster onto a plane, creating a Feature Map (Fig. 4B), in which highly correlated features are mapped nearby each other (see “Feature Interpretation” in Methods for an explanation of individual feature names). In certain cases, these visualizations revealed the nature of the morphological phenotype (e.g., nuclear shape abnormalities distinguishing Cluster 7A PDF), but for others these approaches did not suffice to yield an obvious phenotypic conclusion (e.g., for Cluster 19, Fig. 4A and 4B).

Third, we hypothesized that leveraging the single-cell resolution of image-based profiling might be highly sensitive in enhancing interpretation, particularly for cases where only a subset of cells is distinctive from negative controls. To test this, for each given cluster of genes together with negative controls we identified 20 subpopulations using k-means clustering on single cell data. We calculated the abundance of cells in each of the 20 subpopulations to determine which are over/under-represented relative to controls for the given cluster (corresponding images are shown; Cluster 1B, 2B, … 25B PDFs. For example, the MAPK pathway activators in Cluster 19 show increased prevalence of a subpopulation of cells with strongly asymmetric ER, mitochondria, and Golgi staining, indicating a cell polarization phenotype (Fig. 4C, and Cluster 19B PDF, Categories 1 and 2), for which there is evidence in the literature (Šamaj, Baluška, and Hirt 2004; Elsum, Martin, and Humbert 2013; Godde et al. 2014). This phenotype was not captured by manual inspection nor the first two approaches (e.g., Cluster 19A PDF, panels a2 and b2).

Encouraged by this, we supplemented the morphological map by compiling these and other visualizations into PDF files for each cluster, summarized in Fig. 5 and provided in full as Supplemental Data files for each cluster. We also noticed that certain subpopulations were similar across several clusters (Supp. Fig. 5 shows sample cell images of each such subpopulation); we annotated their enrichment/de-enrichment on the dendrogram (Fig. 3).

**Figure 4:**
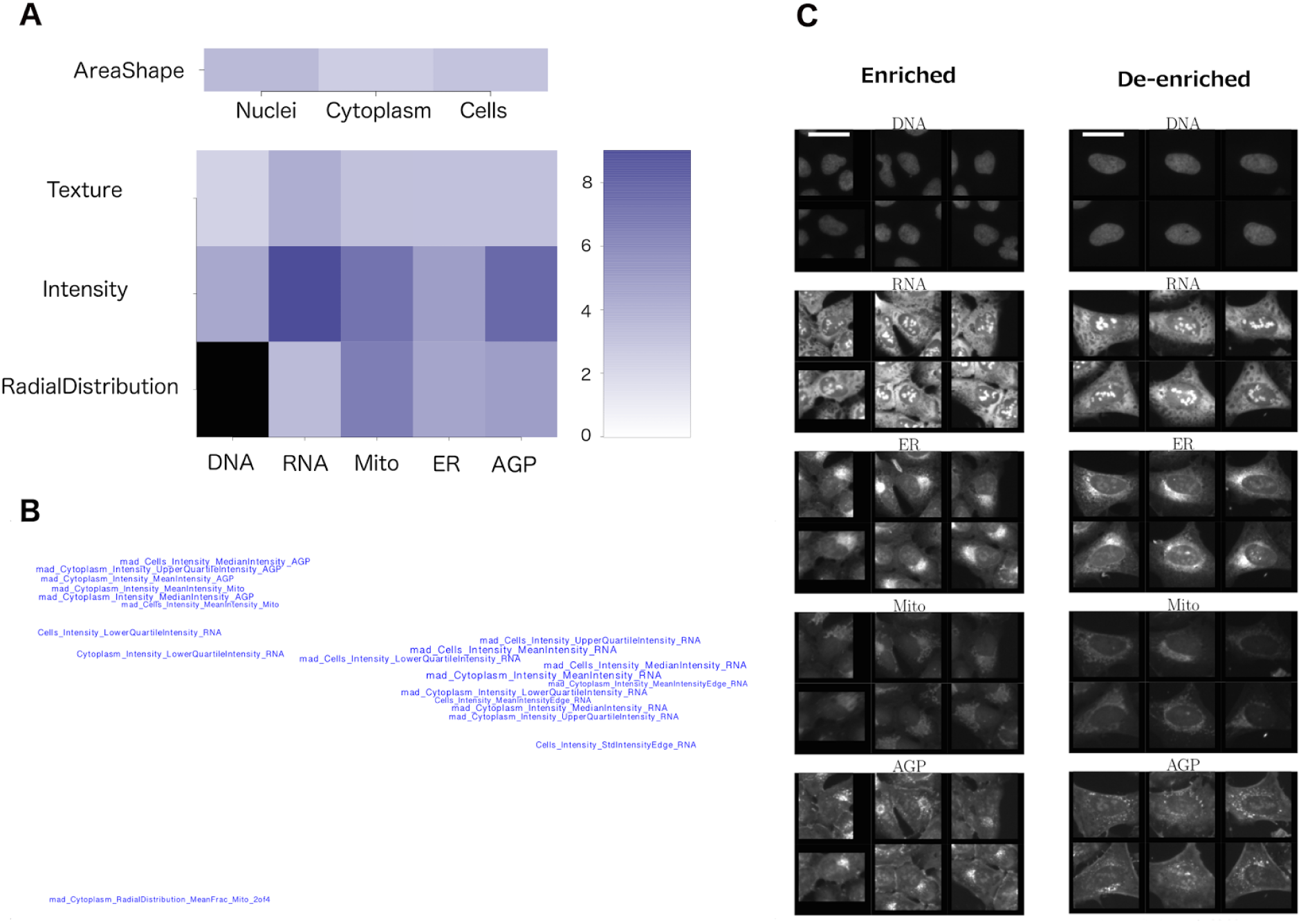
**Visualizations used to interpret morphology of Cluster 19 (for other clusters, see Cluster PDF files). (A) Feature Grid**. RNA and AGP (actin, Golgi, plasma membrane) intensity contribute most to the genes in Cluster 19 (KRAS, RAF1, BRAF, and MOS). Dark blue colors indicate higher median z-score of the relevant measurements for genes in the cluster relative to negative controls. As “RadialDistribution” features do not exist for the DNA channel, it is colored in black. **(B) Feature Map**. The feature names showing the greatest difference between the cluster and negative controls are shown, based on largest absolute value of z-scores (full resolution version is available in Cluster 19A PDF). They are mapped in 2D space such that features that are highly correlated with each other across all genes’ profiles are placed close together and thus can be interpreted together. Blue/red colored names indicate positive/negative sign of the z-score (i.e., blue indicates that the cluster shows higher values than controls). According to this map, the average intensity of AGP, RNA and Mito shows high variation for cells within samples in Cluster 19 (e.g., large mad_Cytoplasm_lntensity_Meanlntensity_AGP, where the prefix “mad” refers to median absolute deviation, a robust form of standard deviation). **(C) Sample images of a subpopulation of cells enriched and de-enriched for all genes in Cluster 19**. Cells with asymmetric organelle distribution are highly over-represented for genes in the cluster, and cells with more even distribution of organelles are less abundant. Scale bars are 39.36 μ*m* long. Pixel intensities are multiplied by 5 for display.

Figure 5: Data and visualizations supporting the morphological map for each cluster. For all 25

**Figure 5:**
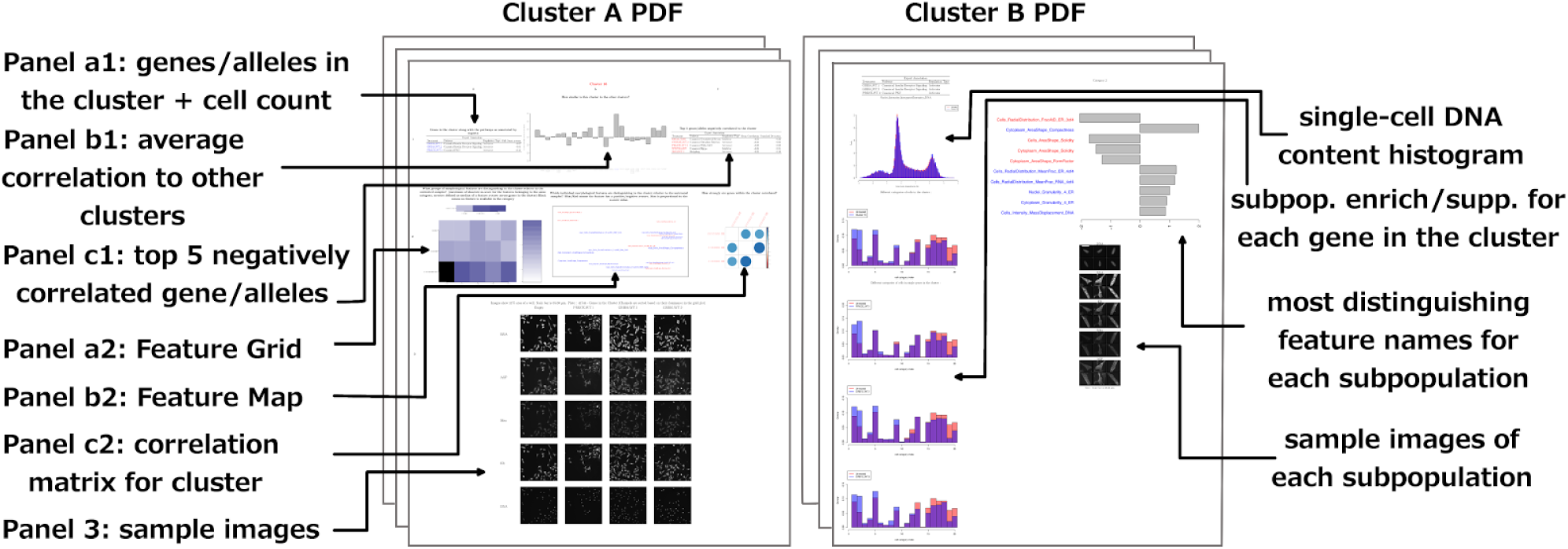
**Data and visualizations supporting the morphological map for each cluster**. **For all 25** clusters, there are two corresponding Supplemental PDF files. **Left: Cluster A PDF files** (e.g., “Cluster_1A.pdf) provide an overview of data about the cluster. **Panel a1** lists the genes/alleles in the cluster as well as expert annotations regarding related pathways and the cell count (as a z-score) for each gene/allele. **Panel b1** contains the average correlation of the cluster to other clusters, indicating uniqueness of the cluster’s morphological phenotype. **Panel c1** lists the top five negatively correlated gene/alleles to the cluster. **Panel a2** shows the Feature Grid summarizing categories of morphological features distinguishing the cluster from the negative control. **Panel b2** shows the Feature Map displaying the names of the top 20 morphological features distinguishing the cluster from the negative control, positioned based on similarity. **Panel c2** shows a correlation matrix for just those genes/alleles in the cluster. **Panel 3** contains sample images of fields of view of cells expressing each gene/allele in the cluster, along with images of the control for comparison. **Right: Cluster B PDF files** contain multiple plots aiming to illustrate the phenotype based on single-cell data, including cell subpopulation enrichment/suppression in the cluster. First, a histogram of single-cell DNA content is shown for all cells from all genes/allele treatments in the cluster, indicating the overall cell cycle distribution. Next, bar plots show (for the cluster overall and for each gene in the cluster) which of 20 subpopulations of cells are enriched and suppressed relative to negative controls. Finally, each subsequent page of the PDF is devoted to the subpopulations whose representation differs from negative controls in a statistically significant way, whether enriched or suppressed (subpopulations which are very small in both the cluster and negative control samples are omitted). For each subpopulation, a bar plot shows the top 10 most-distinguishing feature names (versus negative control cells). Then, sample images are shown of individual representative cells from each subpopulation.

Using these visualizations, we began by interrogating three adjacent and correlating clusters (Clusters 4, 7, and 11) contain wild-type and mutant alleles of CDC42, a gene encoding a Rho family GTPase with diverse roles in cell polarity, morphology, and migration (Melendez, Grogg, and Zheng 2011; Martin 2015). Cluster 4 contains the constitutively active mutant CDC42 Q61L (Nobes and Hall 1999) as well as MAP3K2 and MAP3K9. The highly similar Cluster 7 contains the dominant negative alleles CDC42 T17N (Nobes and Hall 1999) and RAC1 T17N (S. Zhang et al. 1995), a related RAS superfamily member. That activating and inhibiting alleles would yield similar phenotypes when overexpressed is not surprising for CDC42 (Melendez, Grogg, and Zheng 2011). Cluster 7 also contains isoforms and alleles of AKT: specifically, AKT3 and the constitutively active E17K alleles of both AKT1 and AKT3 (M. S. Kim et al. 2008; M. A. Davies et al. 2008). Akt is known to be essential for certain Cdc42-regulated functions (Higuchi et al. 2001) and vice versa (Stengel and Zheng 2012). Finally, the nearby Cluster 11 (which is discussed in more detail later) contains the wild-type form of CDC42 as well as TRAF2, a canonical NF-**κ**B activator; these two are known to interact and share functions in actin remodeling (Marivin et al. 2014). We also note that anti-correlating genes to these clusters (generally in Clusters 13 and 21) are consistent with existing knowledge, including (a) AKT family member AKT1S1 (a Proline rich AKT substrate, PRAS40 (Kovacina et al. 2003; Wiza et al. 2014), Cluster 7A PDF panels b1 and c1) (b) CDK2 (a known target of Akt (Maddika et al. 2008)), (c) PIK3R1 and PTEN in Cluster 21, described previously, which have known interactions with AKT (Cheung and Mills 2016; Hemmings and Restuccia 2015). Thus, all of these connections have previously been identified.

Subpopulation visualization revealed that Clusters 4, 7, and 11 are enriched in cells that are huge and binucleate (Fig. 3, example images shown in Cluster 4B PDF). Genes in all three clusters also show irregularities in DNA content, namely, an enrichment in cells with sub-2N DNA content, a decrease in cells with 2N DNA content, and, for most genes, a decrease in cells with S phase and 4N DNA content, indicating a significant amount of DNA fragmentation and thus apoptosis (DNA histograms in Cluster 4B, 7B, and 11B PDFs). These phenotypes are consistent with these genes’ known role in the cell cycle and cell polarity (Chircop 2014).

As a second test case, we examined Cluster 8, which contains PRKACA (the catalytic subunit α of protein kinase A, PKA) and two of its known substrates: GLI1 (a transcription factor mediating Hedgehog signaling)(Asaoka 2012), and RHOA^Q63L^ (a Ras homolog gene family member)(Lang et al. 1996; Rolli-Derkinderen et al. 2005). The highly similar Cluster 10 contains the wild-type RHOA, as well as ELK1 which is also linked to the Rho GTPase family and PKA (Bachmann et al. 2013; Murai and Treisman 2002).

We investigated the morphological changes causing these genes to cluster. RhoA is a known regulator of cell morphology and cell rounding is a known related phenotype (Oishi et al. 2012). We found that indeed all members of Clusters 8 and 10 significantly induce cell rounding (Supp. Table 7). Although cell count is lower for genes Clusters 8 and 10, the degree varies greatly (from z-score -0.67 to -3.02, Cluster 8A and 10A PDFs, panel a1), ruling out that simple sparseness of cells explains their high similarity in the assay. As well, the overall DNA content distribution of the cell populations appears relatively normal (Cluster 8B and 10B PDFs). Subpopulation extraction provides a satisfying biological explanation for these clusters’ distinctive phenotype: the increased roundness and strong variation in intensity levels (per the Feature Grid) across the population stems from an increased proportion of telophase, anaphase, and apoptotic cells (Fig. 3 and Cluster 8B and 10B PDFs).

We therefore conclude that the morphological map can link related genes to each other and that the morphological data can provide insight into their functions, particularly with the help of subpopulation visualization.

### An unexpected relationship between the Hippo pathway and regulators of NF-**κ**B signaling (Clusters 11, 20, and 22)

We wondered whether novel relationships might emerge from our unbiased classification of gene and allele function based on morphologic profiling. We noticed that the known regulator of NF-**κ**B signaling, TRAF2 (in Cluster 11, together with CDC42) (Grech et al. 2004; Tada et al. 2001), yields a signature strongly anti-correlated to YAP1/WWTR1 (Cluster 20), which encode the transcriptional effectors of the Hippo pathway, YAP (Yes-associated protein) and TAZ (Transcriptional co-activator with a PDZ-domain). The Hippo pathway and NF-**κ**B signaling are critical regulators of cell survival and differentiation, and dysrégulation of these pathways is implicated in a number of cancers (Varelas 2014; Hoesel and Schmid 2013; Tornatore et al. 2012), but we found no evidence in the literature (in particular through BioGRID) of physical interaction between the proteins encoded by Cluster 11 genes and Cluster 20 genes. Confirming our approach, a functional connection between CDC42 (Cluster 11) and YAP1 (Cluster 20) has been identified: deletion of CDC42 phenocopies the loss of YAP1 in kidney-specific conditional knockouts in mice (Reginensi et al. 2013). Still, the NF-**κ**B pathway (and in particular the Cluster 11 member TRAF2), has not been closely tied to YAP and TAZ in human cells (see Discussion).

We first wanted to characterize Clusters 11 and 20 to confirm that relationships within each cluster are supported in the literature. Indeed we found evidence for most of the within-cluster connections. CDC42 and TRAF2 (Cluster 11) physically interact and share functions in actin remodeling (Marivin et al. 2014). As described in a prior section YAP/TAZ (Cluster 20) are known to share functional similarities in the Hippo pathway, being regulated by, and also regulating, cytoskeletal dynamics. Consistent with these known functions, we found that core effector of the Hippo pathway which functions to restrict YAP/TAZ nuclear activity, STK3 (which encodes the Mst2 kinase) (Meng, Moroishi, and Guan 2016), has a morphological signature strongly anti-correlated to YAP1/WWTR1 (Cluster 20A PDF, panel c1). We also note that two clones of another protein known to influence YAP activity, STK11, form Cluster 22 which falls nearby YAP1/WWTR1; a connection between STK11 and YAP has been identified (albeit with opposite directionality, identified via knockdown of STK11 (Mohseni et al. 2014)). Further, YAP1 is among the highest anti-correlating genes to REL (data not shown; REL is a singleton in the dendrogram and thus not in a cluster), whose protein product, c-Rel, has a known connection to TRAF2 (Jin et al. 2015). These results reaffirm that the Cell Painting-based morphological signatures are a useful reporter of biologically meaningful connections among genes in these pathways.

Given the striking inverse correlation between YAP1/WWTR1 and TRAF2, we sought to confirm a negative regulatory relationship between the Hippo and NF-**κ**B pathways by multiple orthogonal methods.

First, we explored the observed inverse morphological impact using the Cell Painting data. The morphological impact of genes in Cluster 11 and 20 is quite strong (median replicate correlation is at the 74th and 81st percentile, and average within-group correlations are 0.66 and 0.73). Subpopulation analysis showed that Cluster 20 (YAP1, WWTR1) is enriched for cells that are slightly large, slightly elongated, and have disjoint, bright mitochondria patterns, whereas Cluster 11 (TRAF2, CDC42) is de-enriched for those subpopulations and instead enriched for binucleate cells, very large cells, and small cells with asymmetric organelles (Fig. 3 and 6A and 6B).

**Figure 6:**
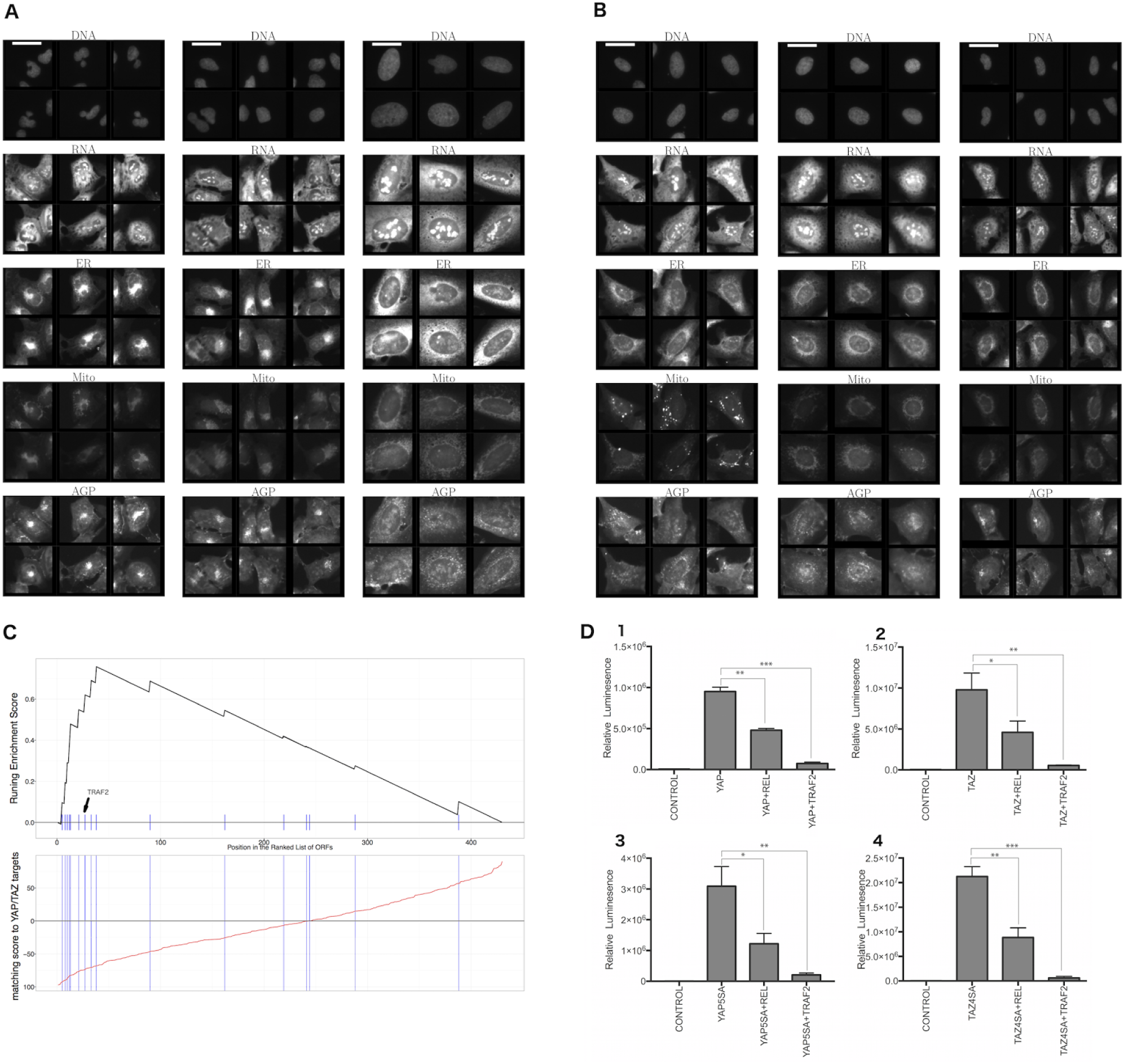
**Morphological and transcriptional cross-talk between the Hippo pathway and regulators of NF-**κ**B signaling. (A)** The TRAF2/CDC42 cluster (Cluster 11) is enriched for bi-nucleate cells, small cells with asymmetric organelles, and huge cells. **(B)** The YAP1/WWTR1 cluster (Cluster 20) is enriched for cells with bright disjoint mitochondria patterns, slightly large cells, and slightly elongated cells. Scale bars are 39.36 μ*m* long. Pixel intensities are multiplied by 5 for display. (C) Gene Set Enrichment Analysis (GSEA) reveals that gene overexpression leading to down-regulation of YAP1 targets (CTGF, CYR61, and BIRC5) are enriched for regulators of the NF-**κ**B pathway (Enrichment Score p-value = 8.19 × 10^-5^)- The horizontal axis gives the index of ORFs sorted based on the average amount of down-regulation of the YAP1 targets. Each blue hash mark on this axis indicates an NF-**κ**B pathway member. The running enrichment score, which can range from -1 to 1, is plotted on the vertical axis and quantifies the accumulation of NF-**κ**B pathways members on the sorted list of ORFs. **(D)** TRAF2 and REL suppress YAP and TAZ transcriptional activity. REL and TRAF2 suppress the ability of wild-type **(D1)** YAP and **(D2)** TAZ to drive the expression of a TEAD-regulated luciferase reporter. Activity of nuclear active mutants of **(D3)** YAP (5SA) and **(D4)** TAZ (4SA) are similarly suppressed. Luciferase reporter activity was measured in HEK293 cells co-transfected with expression constructs as indicated and a TEAD luciferase reporter was used to measure YAP-directed transcription. (* p-value < 0.05, ** p-value = 0.001, *** p-value < 0.0001)

Second, given that YAP/TAZ are transcriptional regulators, we analyzed gene expression data. Using the same constructs as in our Cell Painting experiment, we found an anti-correlated relationship at the mRNA level, consistent with the anti-correlation we had seen in morphological space. To do this, we used Gene Set Enrichment Analysis (Subramanian et al. 2005) and publicly available data, which includes data from four to nine different cell lines at one to four time points (http://lincscloud.org). This analysis revealed that the NF-**κ**B pathway is the pathway most enriched among genes whose overexpression results in down-regulation of known YAP1 targets, CTGF, CYR61, and BIRC5 (Zhao et al. 2008) (adjusted p-value = 2 × 10^-8^, Fig. 6C), with TRAF2 being among the genes contributing to this enrichment (Supp. Table 8). We also saw enrichment of NF-**κ**B pathway members when testing a data-driven set of targets of YAP1/TAZ (Supp. Fig. 7, see Methods). In the inverse analysis, genes that alter the levels of TRAF2/REL common targets are weakly enriched in Hippo pathway members (Supp. Fig. 8, see Methods). This is consistent with the hypothesis that NF-**κ**B members can downregulate YAP/TAZ targets but not strongly vice versa.

Finally, we more directly confirmed negative crosstalk between NF-**κ**B effectors and YAP/TAZ using a synthetic TEAD luciferase reporter that is YAP/TAZ responsive (Dupont et al. 2011). Importantly, these confirmatory experiments used different cellular contexts and perturbation constructs versus the original Cell Painting data. Co-expression of the NF-**κ**B pathway effectors TRAF2 or C-REL with YAP or TAZ led to significantly lower reporter activity than expression of YAP or TAZ alone (Fig. 6D1 and 6D2). Intriguingly, mutants of YAP or TAZ that are insensitive to negative regulation by the Hippo pathway (YAP-5SA and TAZ-4SA; (Zhao et al. 2008)) remained sensitive to suppression of transcriptional activity by TRAF2 and C-REL, indicating that the negative relationship we identified may be independent of canonical upstream Hippo pathway signals (Fig. 6D3 and 6D4).

## Discussion

We conclude that connections among genes can be profitably analyzed using morphological profiling of overexpressed genes via the Cell Painting assay. In a single inexpensive experiment, we were able to rediscover a remarkable number of known biological connections among the genes tested. Further, we found that morphological data from the Cell Painting assay, together with novel subpopulation visualization methods, can be used to flesh out the functionality of particular genes and/or clusters of interest.

By adopting a two-pronged approach, merging this Cell Painting morphological analysis with transcriptional data, we were able to identify an unexpected relationship in human cells between two major signaling pathways, Hippo and NF-**κ**B, both under intense study recently for their involvement in cancer. Through validation of these clustered genes, we have identified that YAP/TAZ-directed transcription is negatively regulated by NF-**κ**B pathway effectors and our data suggests a novel regulatory mechanism that is independent of upstream Hippo kinases.

To date, there has been little evidence of the intersection between these important signaling pathways. Recent work examining osteoclast-osteoblast differentiation has suggested that Hippo pathway kinases, such as Mst2, may affect the NF-**κ**B pathway through phosphorylation of IkB proteins, thereby promoting nuclear translocation of NF-**κ**B transcription factors (Lee et al. 2015). TAZ was found to be a direct target of NF-**κ**B transcription factors and its expression is regulated via NF-**κ**B signaling (Cho et al. 2010). Our work, however, supports a possible additional mode of interaction, whereby regulators of NF-**κ**B signaling directly regulate the function of Yap and Taz as transcriptional co-factors. Recent work has demonstrated, in *Drosophila,* that NF-**κ**B activation via Toll receptor signaling negatively regulates the transcriptional activity of Yorkie, the homolog of YAP/TAZ, through activation of canonical hippo pathway kinases (B. Liu et al. 2016). The work described here identifies, for the first time in a mammalian system, that a negative regulatory relationship exists between NF-**κ**B activation and YAP/TAZ transcriptional function. Furthermore, we have identified that this regulation of YAP/TAZ occurs in a manner that is independent of Hippo pathway-mediated phosphorylation events on YAP/TAZ, suggesting a more direct relationship between NF-**κ**B and YAP/TAZ signaling.

In this work, we tested quantitatively and explored qualitatively the connections among genes revealed by morphological profiling. Our underlying hypothesis was that functionally similar genes would generally yield morphologically similar cells when overexpressed, and indeed we found this to be the case. Still, some discussion of this point is warranted. Most commonly, gene overexpression will result in activation of the corresponding pathway via amplification of the endogenous gene’s function. However, it is important to note that the profiling strategy to discover functional relationships does not assume or require this. For example, overexpression could also disrupt a protein complex, producing a trans-dominant negative effect that results in precisely the opposite phenotypic effect (Veitia 2007). In still other cases, overexpression of a particular gene may not affect any of the normal functions of the gene (producing a false negative signal), or trigger a stress response (yielding a confounded profile), or produce a complicated response, due to feedback loops. Further, artifactual phenotypes could be seen, e.g., if overexpression yields a non-physiological interaction among proteins or toxic aggregates. Nevertheless, despite these caveats and complications, our results indicate that valuable information could be gleaned from the similarity and dissimilarity of the morphological perturbations induced by gene overexpression. Using overexpression avoids the complications of RNAi off-target effects (often due to seed effects), which were far more prevalent (impacting 90% of constructs in our recent study (Singh et al. 2015)).

In addition to functionally annotating genes, as demonstrated here, one particularly appealing application enables personalized medicine: it should be feasible to use morphological profiling to predict the functional impact of various disease alleles, particularly rare variants of unknown significance. This has recently been successful using mRNA profiles (Berger et al. 2016). Thus, an even more exciting prospect would be to combine mRNA profiles with morphological profiles to better predict groups of alleles of similar mechanism, and ultimately to predict effective therapeutics for each group of corresponding patients.

We make all raw images, extracted cellular features, calculated profiles, and interpretive visualizations publicly available, providing an initial morphological map for several major signaling pathways, including several unexplored connections among genes for further study (see Supp. Data files). Expanding this map to full genome scale could prove an enormously fruitful resource.

## Materials and Methods

### cDNA constructs used for expression

The Reference Set of human cDNA clones utilized here has been previously described (E. Kim et al. 2016); ~90% of these constructs induce expression of the intended gene greater than 2 standard deviations above the control mean. Briefly, wild-type ORF constructs were obtained as Entry clones from the human ORFeome library version 8.1 (http://horfdb.dfci.harvard.edul with additional templates generously provided by collaborating laboratories, and cloned into the pDONR223 Gateway Entry vector. In addition, here, to maximize coverage of cellular pathways, we included additional clones with minimal sequence deviations from the intended templates. Sanger sequencing of Entry clones verified the intended transcripts and, if applicable, the intended mutation. Entry constructs and associated sequencing data will be publicly available via www.addaene.ora and may also be available via members of the ORFeome Collaboration (http://www.orfeomecollaboration.org/), including the Dana-Farber/Harvard Cancer Center (DF/HCC) DNA Resource Core DNA Repository (http://www.dfhcc.harvard.edu/core-facilities/dna-resource/) and the DNASU Plasmid Repository at ASU Biodesign Institute (http://dnasu.asu.edu/DNASU/Home.jsp). Clone requests must include the unique clone identifier numbers provided in the last column of Supp. Table 1 (e.g. ccsbBroadEn_12345 as an example for a specific entry clone and ccsbBroad304_12345 as an example for a specific expression clone). ORFs were transferred to the pLX304 lentiviral expression vector (Yang et al. 2011) by LR (attLx attR) recombination.

For simplicity, throughout this paper “wild-type” refers to ORFs found in the original collection without a particular known mutation intentionally engineered. Due to natural human variation, and occasional cloning artifacts, there are often non-identical matches of such constructs to reference sequence; these differences are fully documented for each construct and sequence data will be publicly available through AddGene, in addition to the sequencing data for the original Entry clones for the genome-scale library(Yang et al. 2011).

### Lentiviral transduction for morphological profiling

We followed our previously described protocol (E. Kim et al. 2016; Berger et al. 2016) except for durations of some steps. Briefly, cells were plated in 384-well plates and transduced with lentiviral particles carrying ORF constructs the next day. Viral particles were removed 18-24 hr post-infection and cells cultured for 48 hr until staining and imaging (72 hr total post-transduction).

### Cell staining and imaging

The Cell Painting assay followed our previously published protocol (Bray et al. 2016). Briefly, eight different cell components and organelles were stained with fluorescent dyes: nucleus (Hoechst 33342), endoplasmic reticulum (concanavalin A/AlexaFluor488 conjugate), nucleoli and cytoplasmic RNA (SYT014 green fluorescent nucleic acid stain), Golgi apparatus and plasma membrane (wheat germ agglutinin/AlexaFluor594 conjugate, WGA), F-actin (phalloidin/AlexaFluor594 conjugate) and mitochondria (MitoTracker Deep Red). WGA and MitoTracker were added to living cells, with the remaining stains carried out after cell fixation with 3.2% formaldehyde. Images from five fluorescent channels were captured at 20x magnification on an ImageXpress Micro epifluorescent microscope (Molecular Devices): DAPI (387/447 nm), GFP (472/520 nm), Cy3 (531/593 nm), Texas Red (562/624 nm), Cy5 (628/692 nm). Nine sites per well were acquired, with laser based autofocus using the DAPI channel at the first site of each well.

### Image processing and feature extraction

The workflow for image processing and cellular feature extraction has been described elsewhere (Bray et al. 2016), but we describe it briefly here. CellProfiler (Carpenter et al. 2006) software version 2.1.0 was used to correct the image channels for uneven illumination, and identify, segment, and measure the cells. An image quality workflow (Bray et al. 2012) was applied to exclude saturated and/or out-of focus wells; six wells containing blurry images were excluded, retaining 1,914 plate/well combinations in the experiment. Cellular morphological, intensity, textural and adjacency statistics were then measured for the cell, nuclei and cytoplasmic sub-compartments. The 1,402 cellular features thus extracted were normalized as follows: For each feature, the median and median absolute deviation were calculated across all untreated cells within a plate; feature values for all the cells in the plate were then normalized by subtracting the median and dividing by the median absolute deviation (MAD) times 1.4826 (Chung et al. 2008). Features having MAD = 0 in any plate were excluded, retaining 1,384 features in all.

### Profiling and data preprocessing

The code repository for the profiling and all the subsequent analysis will be made publicly availabe at https://aithub.com/carpenterlab/2016 rohban submitt. We will next explain details of each analysis step implemented in the code. Single cell measurements in each well and plate position are summarized into the profiles by taking their median and median absolute deviation (abbreviated as “MAD” or “mad” in some tables) over all the cells. Although this method does not explicitly capture population heterogeneity, no alternate method has yet been proven more effective (Ljosa et al. 2013). We also include the cell count in a sample as an additional feature. This results in a vector of 2,769 elements describing the summarized morphology of cells in a sample. We then use the median polishing algorithm immediately after obtaining the summarized profiles, to remove and correct for any plate position artifacts. For each feature, the algorithm de-trends the rows, i.e. by subtracting the row median from the corresponding feature of each profile in that particular row. Next, it de-trends the columns in a similar way using column medians. The row and column de-trending is repeated until convergence is reached in all the features. For the rest of the analysis we considered only the constructs which have more than 99% sequence identity to both the intended protein and gene transcript, to avoid testing uncharacterized mutations/truncations.

Not all of the morphological features contain useful reproducible information. We first filter out features for which their replicate correlation across all samples (except the negative controls) is less than 0.30, retaining 2,200 features. Subsequently, a feature selection method is used (Fischer et al. 2015). Briefly, starting with features (measurements) that we identify as essential, a new feature that contributes the most information with respect to those that have been chosen, is added to the set. The contribution of each feature to the already-selected features is measured by the replicate correlation of the residue when the feature is regressed on the already selected features. This is repeated until the incremental information added drops below a threshold. The original method proposed in (Fischer et al. 2015) overfits in its regression step when the original data is very high dimensional. As a remedy, in the regression step we only use features that have a Pearson correlation of more than 0.50 with the selected features thus far. This prevents overfitting of regression when the dimensionality of selected features grows. We stop feature selection when the maximum replicate correlation of residue is less than 0.30.

The feature selection method greatly removes redundancy, but because of the non-optimal “greedy” strategy, some redundancy remains. Principal component analysis is then applied to keep 99% of variance in data, resulting in 158 principal components being selected.

### Feature interpretation

The features measured using CellProfiler follow a standard naming convention. Each feature name is made up of several tokens separated by underscores, in the following order:

- **Prefix** which could be either empty or “mad”. This means that the feature is calculated either by taking median (no prefix) or median absolute deviation (“mad” prefix) of the relevant measurement over all the cells in a sample.
- **Cellular compartment** in which the measurement related to the feature is made, i.e., “Cells”, “Cytoplasm”, or “Nuclei”. Note that features labeled “Nuclei” are based on segmentation of nuclei using Hoechst staining, “Cells” are based on segmentation of the cell edges using the RNA channel, and “Cytoplasm” is the subtraction of the aforementioned compartments.
- **Measurement type,** which can be either “Intensity”, “Texture”, “RadialDistribution”, “AreaShape”, “Correlation_Correlation”, “Granularity”, and “Neighbors”. Note that “Correlation_Correlation” measures, within a cellular compartment, the correlation between gray level intensities of corresponding pixel pairs across two channels (specified in the next tokens in the feature name). Note also that the relative positioning of a cell is measured in the “Neighbors” category.
- **Name(s) of channels** in which the measurement is made, if appropriate (omitted for AreaShape and Neighbors).
- **Feature name**. The precise measurement name appears at the end. A description of each metric can be found in the Cell Profiler manual (http://d1zymp9ayga15t.cloudfront.net/CPmanual/index.html)

### Identifying ORF constructs that are distinguishable from negative controls

Our method to identify which genes produce a discernable profile involves first normalizing each profile to the negative controls, such that a treatment’s median replicate correlation becomes a surrogate for phenotype strength. The cutoff for “discernible” is set based on the top 5th percentile of a null distribution. The null distribution is defined based on the correlations between non-replicates (that is, different constructs) in the experiment. Treatments whose replicate correlations are greater than the 95th percentile of the null distribution are considered as “hits” that have a morphological phenotype that is highly reproducible (Fig. 2A).

At this point, for strong treatments, all profiles of the replicates are collapsed by taking the average of individual features. 110 out of the 112 selected ORFs were significantly different from the untreated profiles in the feature space. That is, their average Euclidean distances to the untreated profiles were higher than 95th percentile of untreated profile distances to themselves. This shows these two alternative notions of phenotype strength-replicate reproducibility and distance to negative control-are consistent. We restrict all the remaining analyses to the 110 ORFs.

### Comparison of morphological connections between genes to protein-protein interaction data and pathway annotations

In this analysis, mutant alleles were removed and we considered only one wild-type allele for each gene with a detectable phenotype, retaining 73 genes. We calculated a threshold to identify significantly correlated gene pairs. We picked the threshold to minimize the probability of error in classifying wild-type clone pairs versus different-gene pairs. To do so, we found the value at which the probability density functions of the two groups intersect (Supp. Fig. 6); this value (here, 0.43) can be proved to have the desired property (Duda, Hart, and Stork 2012). This approach results in about 5% of the gene pairs being categorized as highly correlated. We next formed a 2 by 2 contingency table, where the rows correspond to two groups of gene pairs, determined by whether they have high profile correlation or not. Similarly, the columns also correspond to two groups of gene pairs, determined by whether the corresponding proteins have been reported to interact in BioGRID (or alternatively have been annotated to be in the same pathway; Supp. Table 3 and 4). This table was then used to perform a one-tailed Fisher’s exact test.

### Creation of a dendrogram relating genes to each other, and aggiomerative clustering by cutting the dendrogram

A dendrogram was created based on the Pearson correlation distance and average linkage, using the hclust function in R (Fig. 3).

Gene clusters were formed by cutting the dendrogram at a fixed correlation level, 0.522, which was chosen using a stability-based measure. The measure is defined as follows: the local clustering stability is measured for a range of candidate cutoffs, from 0.43 (used earlier to test consistency to protein interaction data) to 0.70. The point with highest stability was chosen (Supp. Fig. 4), and the stability measure was defined as the proportion of treatments whose clusters do not change if the cutoff is slightly changed by a small amount, ε = .002 ■

### Subpopulation extraction

In order to extract cell categories (subpopulations) and subpopulation enrichment laid over the dendrogram in Fig. 3, we applied k-means clustering on the normalized single cell data for each gene cluster and the control. Data normalization was carried out on a plate-wise basis by z-scoring each feature using the control samples as reference. In order to avoid curse of dimensionality, we restricted the dataset to the features obtained from the feature selection step mentioned earlier. We set k=20 to be the number of subpopulations. The algorithm was run for at most 5000 iterations. Each cell was assigned to the subpopulation for which it has the shortest Euclidean distance to its center. Then, the number of cells belonging to each cell subpopulation was counted and the proportion in each subpopulation for genes in the cluster was compared against that of the control. If the change in proportion of a cell category was consistent across the genes in the cluster, the cell category is shown in the supplemental PDFs. To quantify this consistency, we used the inverse coefficient of variation of the change in a category proportion. If this quantity exceeded 1, we called the change consistent and included the corresponding cell category in the PDFs. Images of cells which have highest similarity to the category center in the feature space are then used to interpret and give name to each cell category (Supp. Fig. 5)

### Identifying targets of a gene using a data-driven approach

For this purpose, we used a replicate of the original experiment but with L1000 gene-expression readouts, which is provided in the supplemental data; i.e. cell line, time point, and ORF constructs are the same. This data is different from the data used in creating GSEA plots, which entails multiple cell lines and time points. The mRNA levels are all normalized with respect to the negative control. For each replicate of the overexpression construct, we sort the expression levels of landmark genes and take the list of top and bottom 50 landmark genes. Then, to find targets of the gene related to the construct, we find the landmark genes among this list which has shown up at least in p% of replicates/clones of the gene. In particular, we set p to 33% for YAP1, 50% for WWTR1, TRAF2, and REL. Then, we simply take the intersection of predicted targets of YAP1 and WWTR1 (and similarly TRAF2 and REL, separately) to get their common targets. These targets are then used to produce Supp. Fig. 7 and 8.

### Gene Set Enrichment Analysis

In order to produce Fig. 6C, we specified the three known targets of YAP/WWTR1 (CYR61, CTGF, and BIRC5) and queried for ORFs resulting in down-regulation of these genes. This scores each ORF (out of the 430 in the dataset) based on the observed change in mRNA level of the specified YAP/WWTR1 targets, across between four to nine different cell lines and between one to four time points. For each ORF, we then sought the summarized score which takes the mean of 4 largest scores across time point/cell line combinations. Finally, the ORFs were sorted based on the summarized score, and top 30 ORFs were tested for enrichment in different pathways (Supp. Table 8). We used the “clusterProfiler” package in R and the KEGG pathway enrichment analysis implemented in it for creating the GSEA plot (Yu et al. 2012).

### Luciferase Reporter assay

Wild-type and mutant sequences of WWTR1 (TAZ) (4SA: S66A, S89A, S117A, and S311A) and YAP1 (5SA: S61A, S109A, S127A, S164A, and S397A) were previously generated and cloned into the pCMV5 backbone; these constructs are distinct from those used in the original Cell Painting data set. TRAF2 and REL were cloned from the original constructs (using Broad ID# ccsbBroadEn_01710 and ID# ccsbBroadEn_11094, respectively), into pCMV5 expression vectors. These were sequenced and confirmed to BLAST against the appropriate Broad clone ID. The empty pCMV5 backbone was used as the control condition. The Tead luciferase reporter construct, 8xGTIIC-luciferase was a gift from Stefano Piccolo (Addgene plasmid # 34615).

HEK293 cells were transfected using Turbofect (ThermoFisher Scientific) according to manufacturer’s protocol. All cells were co-transfected with a ε-galactosidase reporter plasmid (pCMV-LacZ from Clontech) as a transfection control. Cells were lysed 48 hours following transfection. Lysates were mixed with firefly luciferase (Promega) according to the manufacturer’s protocol and luminescence was measured using a luminometer (BioTek). Lysates were mixed with o-nitrophenyl-ε-D-galactoside (ONPG) and ε-galactosidase expression was determined spectrophotometrically by measurement of absorbance at 405nm following ONPG cleavage. All luciferase readings were normalized to ε-galactosidase expression for the sample. Statistical analysis was conducted using a two tailed unpaired Student’s t test. The data shown in Fig. 6D are from triplicate samples within a single experiment and is representative of replicate experiments.

## Acknowledgements

The authors gratefully acknowledge contributions from members of the Carpenter laboratory, especially Steven A. Moore. Funding for this work was provided by the National Science Foundation (NSF CAREER DBI 1148823 to AEC), a BroadNextIO grant from the Broad Institute, and the Slim Initiative for Genomic Medicine, a project funded by the Carlos Slim Foundation in Mexico.

## Supplementary Data/Figures

### Supplemental data

We will upload the L1000 gene expression data to GEO

(https://www.ncbi.nlm.nih.gov/geo/query/acc.cgi?acc=GSE70138) and the image data along with the extracted morphological features at the per-cell level to the Image Data Repository (http://idr-demo.openmicroscopy.org/about).

As supplemental data files, we provide:

- Two PDF-formatted files (A and B) containing details about each of the 25 clusters (50 total).
- Files containing the per-cell and per-well features extracted by CellProfiler.
- The CellProfiler pipeline used to quantify the images (.cppipe format).

**Supp. Figure 1:**
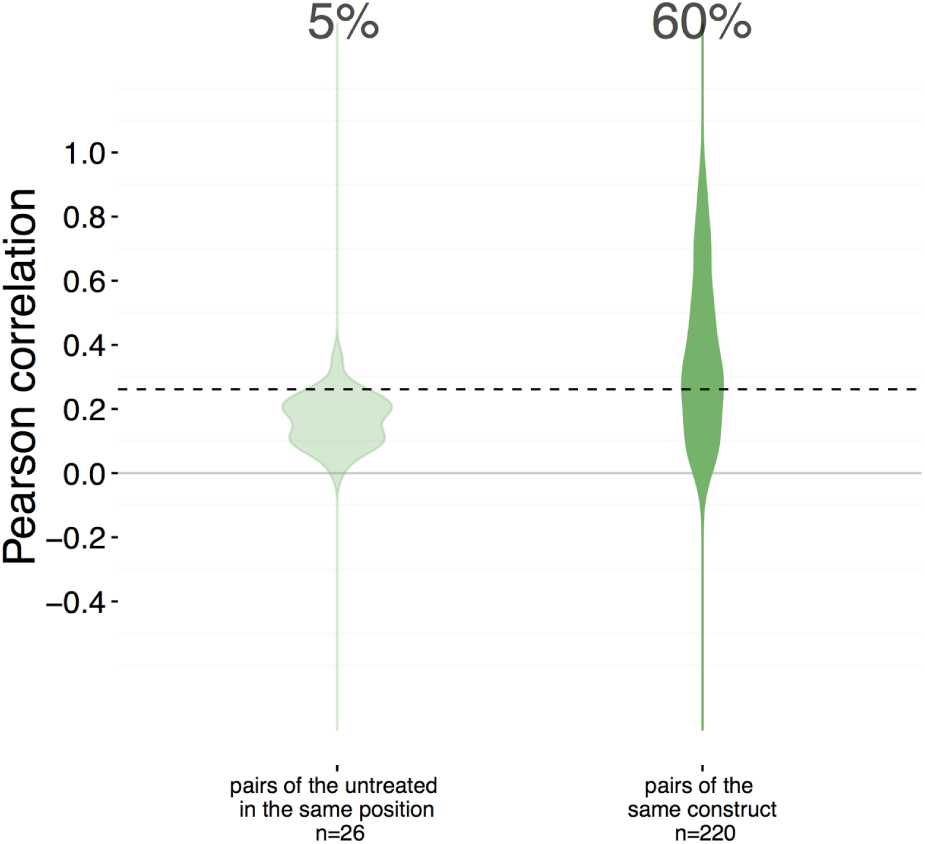
**Position artifacts do not contribute to the hit rate seen in the experiment**. We were concerned that position artifacts may result in overestimating the replicate correlations because replicates of the same treatment are assigned to the same well location across different plates in the experiment (that is, it was infeasible to scramble well locations). We ruled out this possibility by taking an alternative pessimistic null distribution which takes well position into account. In contrast to Figure 2A, which shows a 51% hit rate, a more pessimistic alternative null distribution is shown here (left), calculated based on the replicate correlation of pairs of negative controls in the same position only. We consider this less reliable because the number of such pairs is small (26) and we excluded edge wells; nevertheless the hit rate increases slightly, to 60%.

**Supp. Figure 2:**
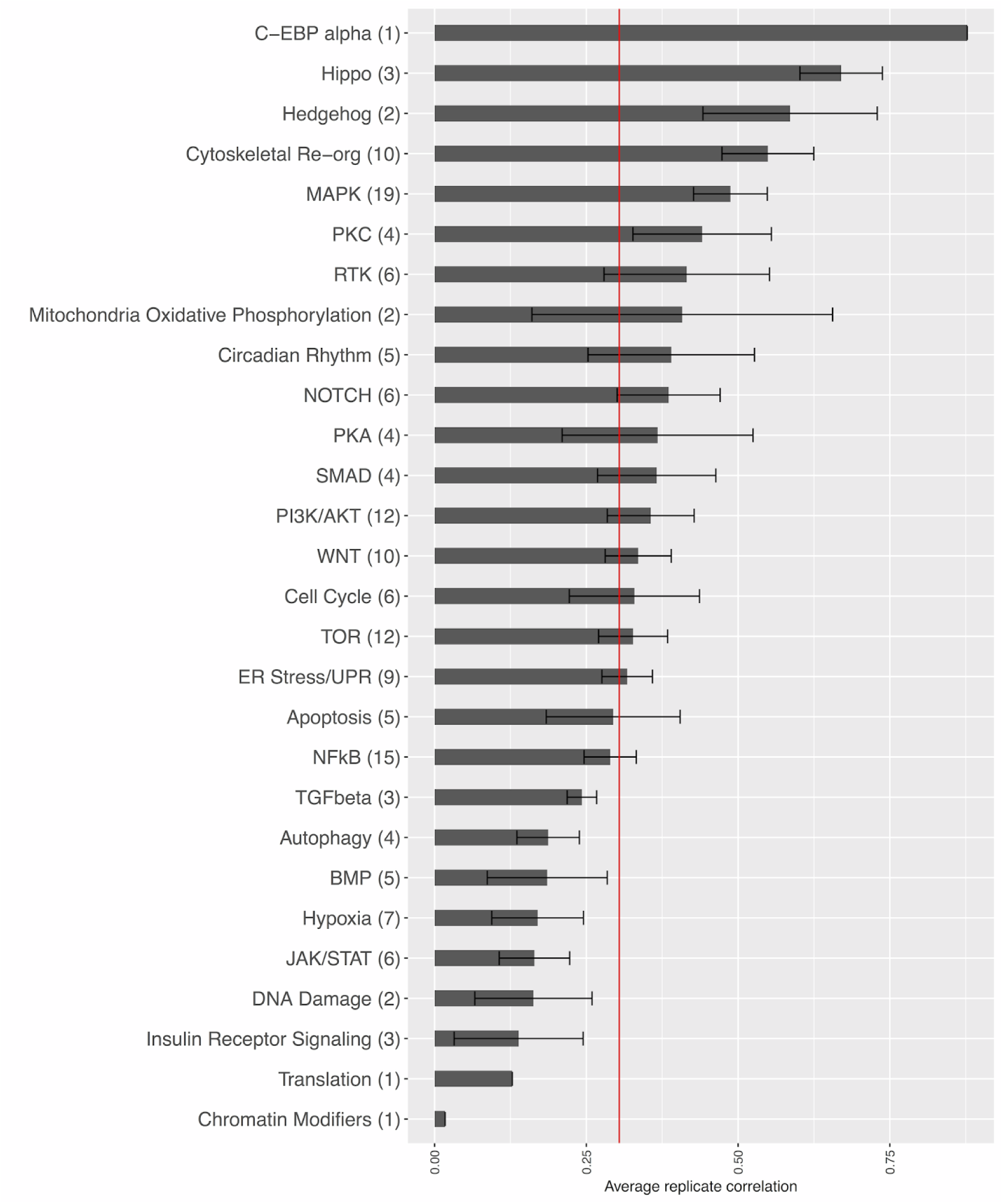
**Strength of morphological phenotypes, according to annotated pathway**. Morphological phenotype strength is calculated as the average replicate correlation for genes that experts manually annotated genes as belonging to each pathway. The number of genes tested in each category is shown in parentheses after the pathway name (two wild-type clones of the same gene are only counted once, but a mutant allele is counted separately). The red line shows the threshold beyond which an individual gene’s profile would be considered to yield a distinguishable phenotype. Error bars indicate the deviation in the replicate correlation among the genes associated with the pathway.

**Supp. Figure 3:**
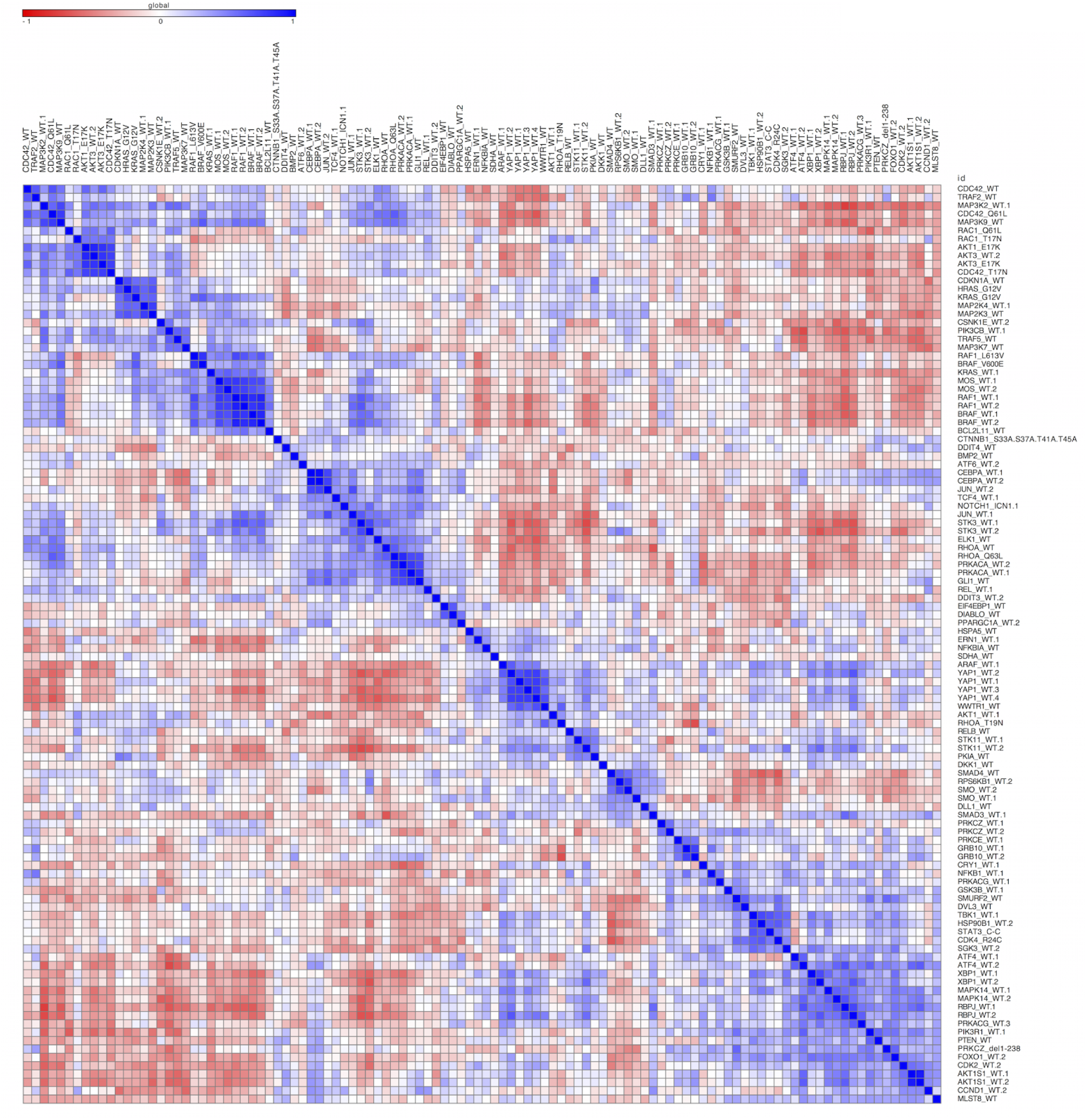
**Correlation among the 110 genes/alleles with a detectable morphological phenotype**. The rows and columns are ordered based on a hierarchical clustering algorithm such that each blue submatrix on the diagonal shows a cluster of genes resulting in similar phenotypes. The scale bar depicts Pearson correlation.

**Supp. Figure 4:**
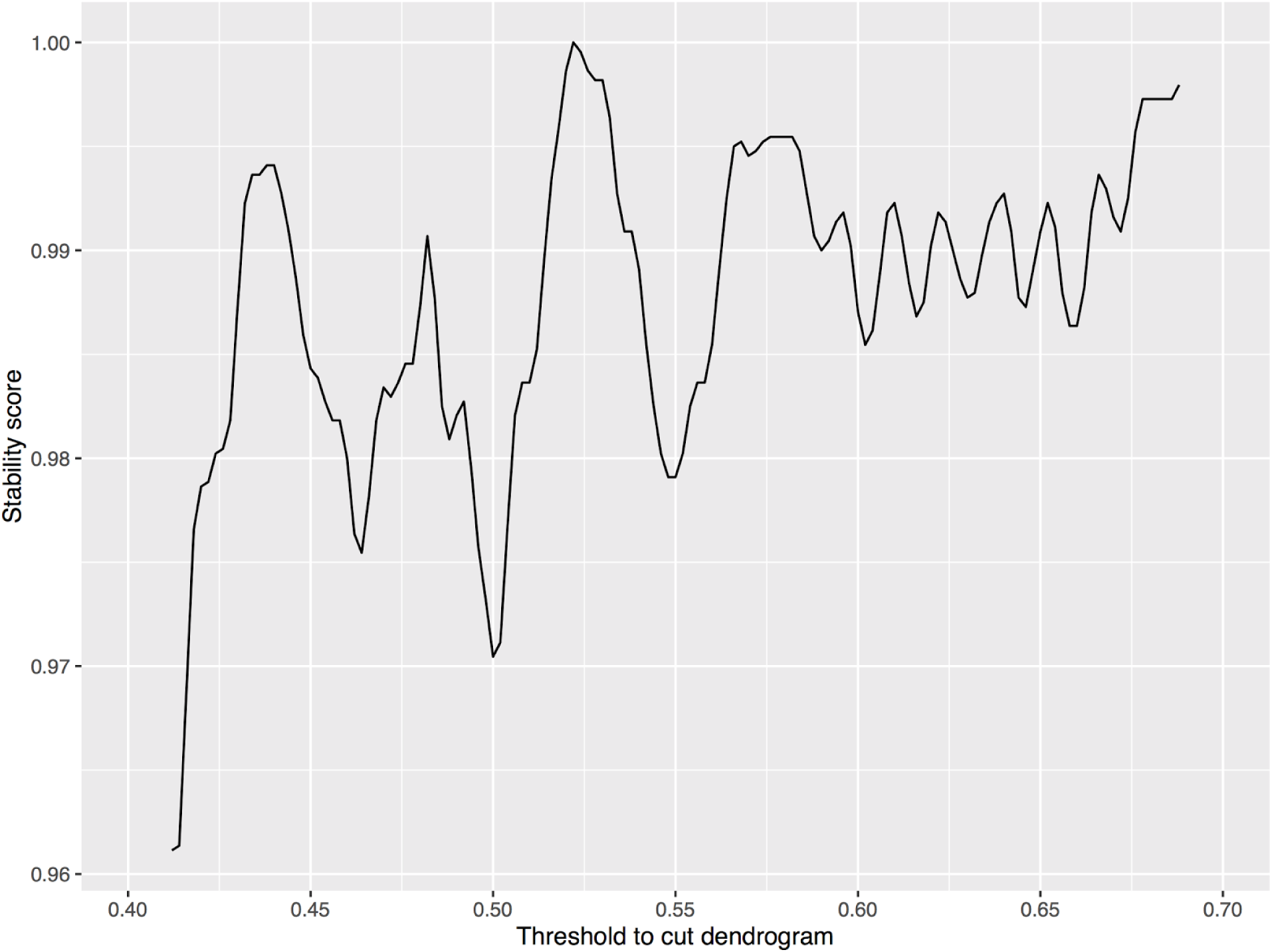
**Smoothed stability score across different cutoffs, in order to choose a threshold for cutting the dendrogram to form clusters**. The maximum occurs at threshold = 0.522. Smoothing is done by taking the moving average of order 0.02. The stability score is defined as the proportion of treatments whose clusters are not affected if the cutoff is increased or decreased by a small amount (ε = .002).

**Supp. Figure 5:**
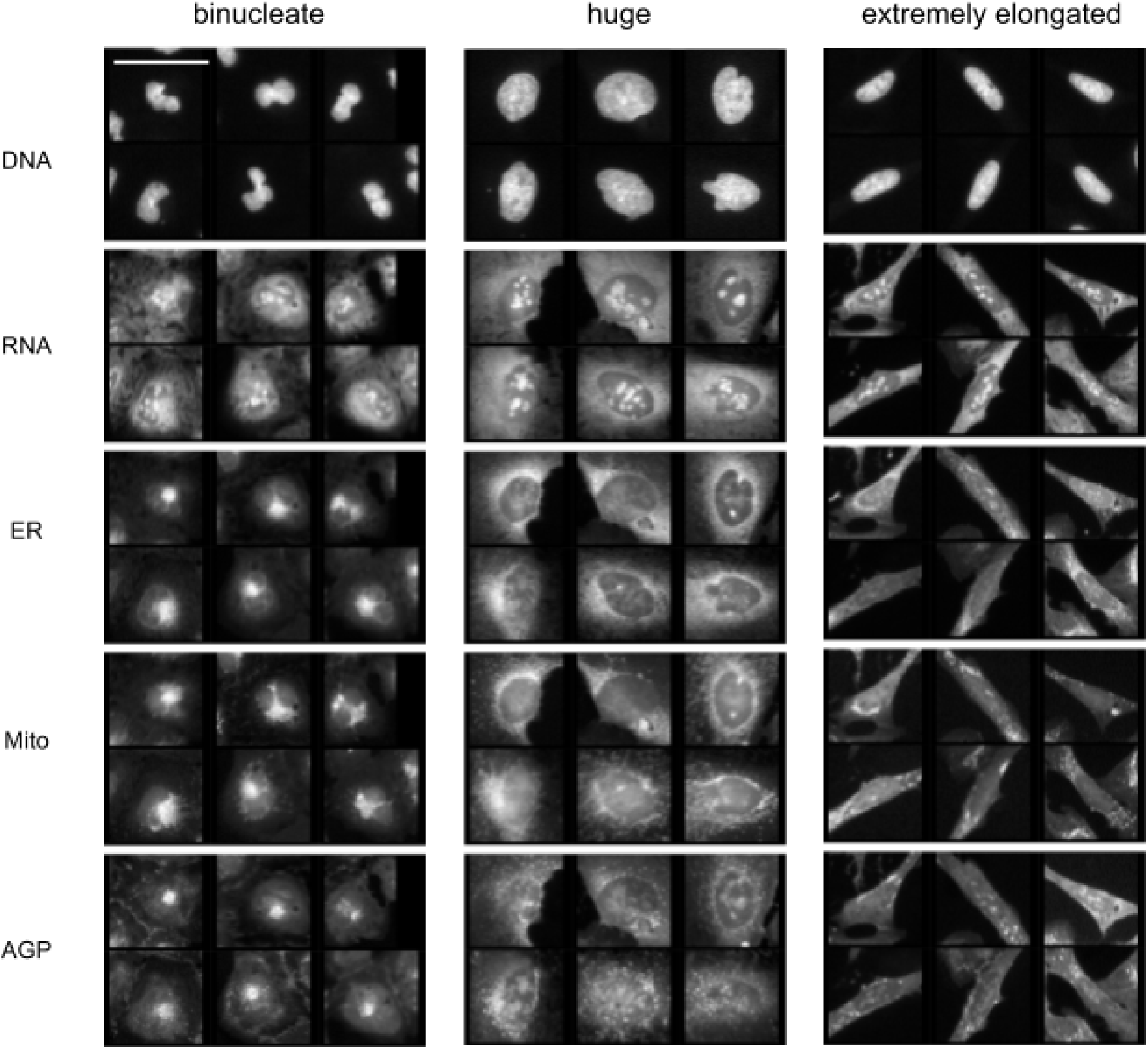
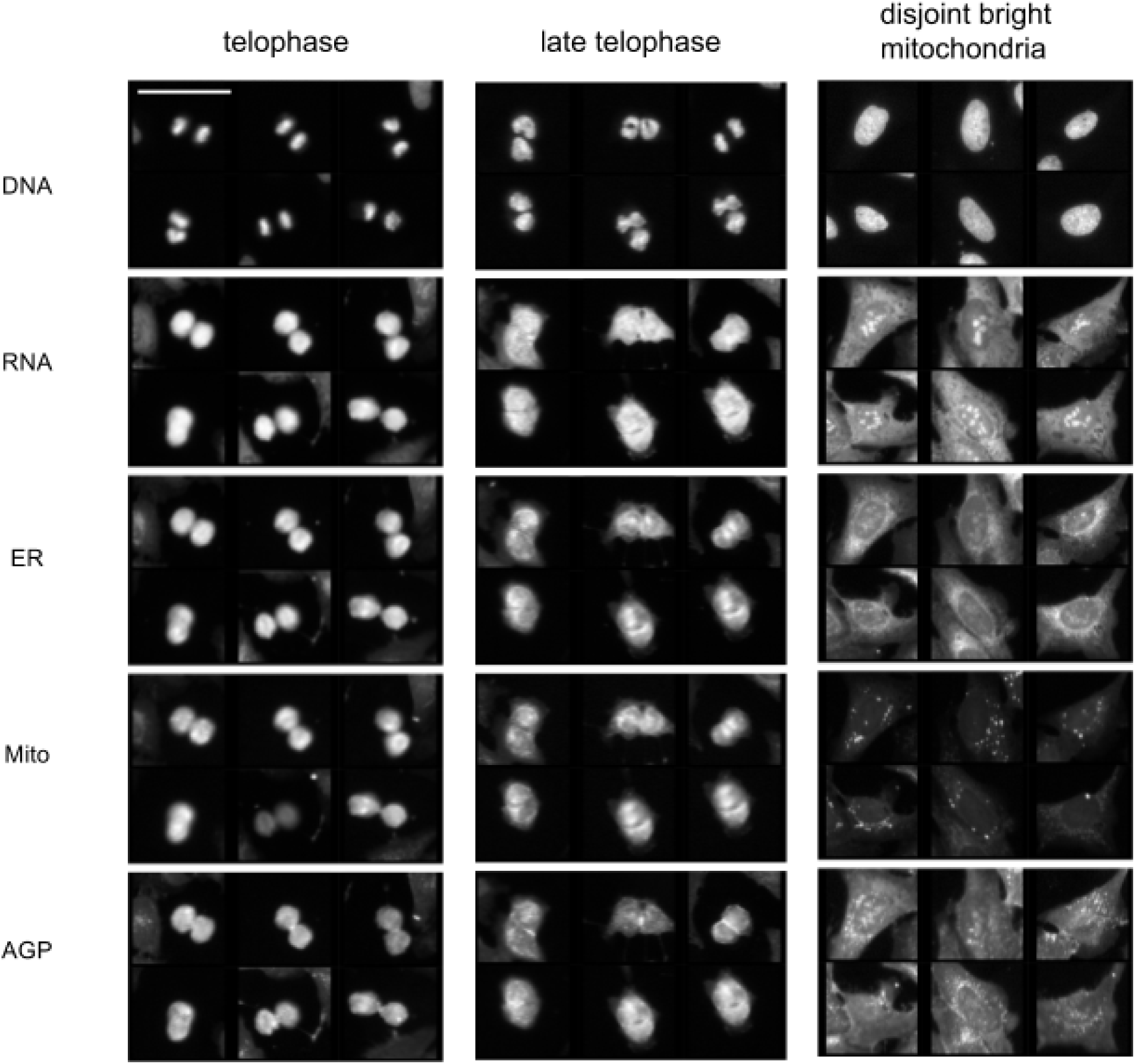
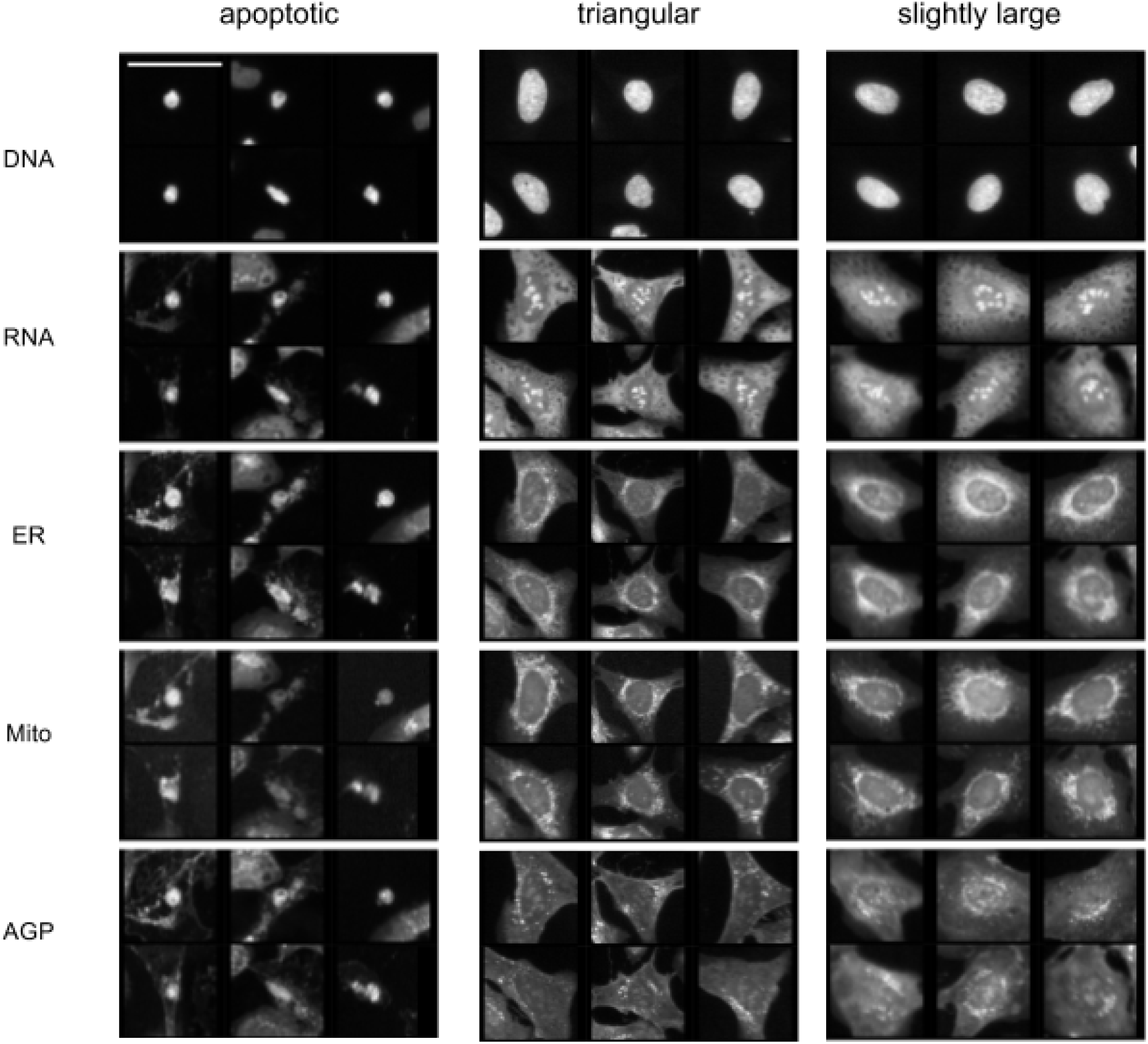
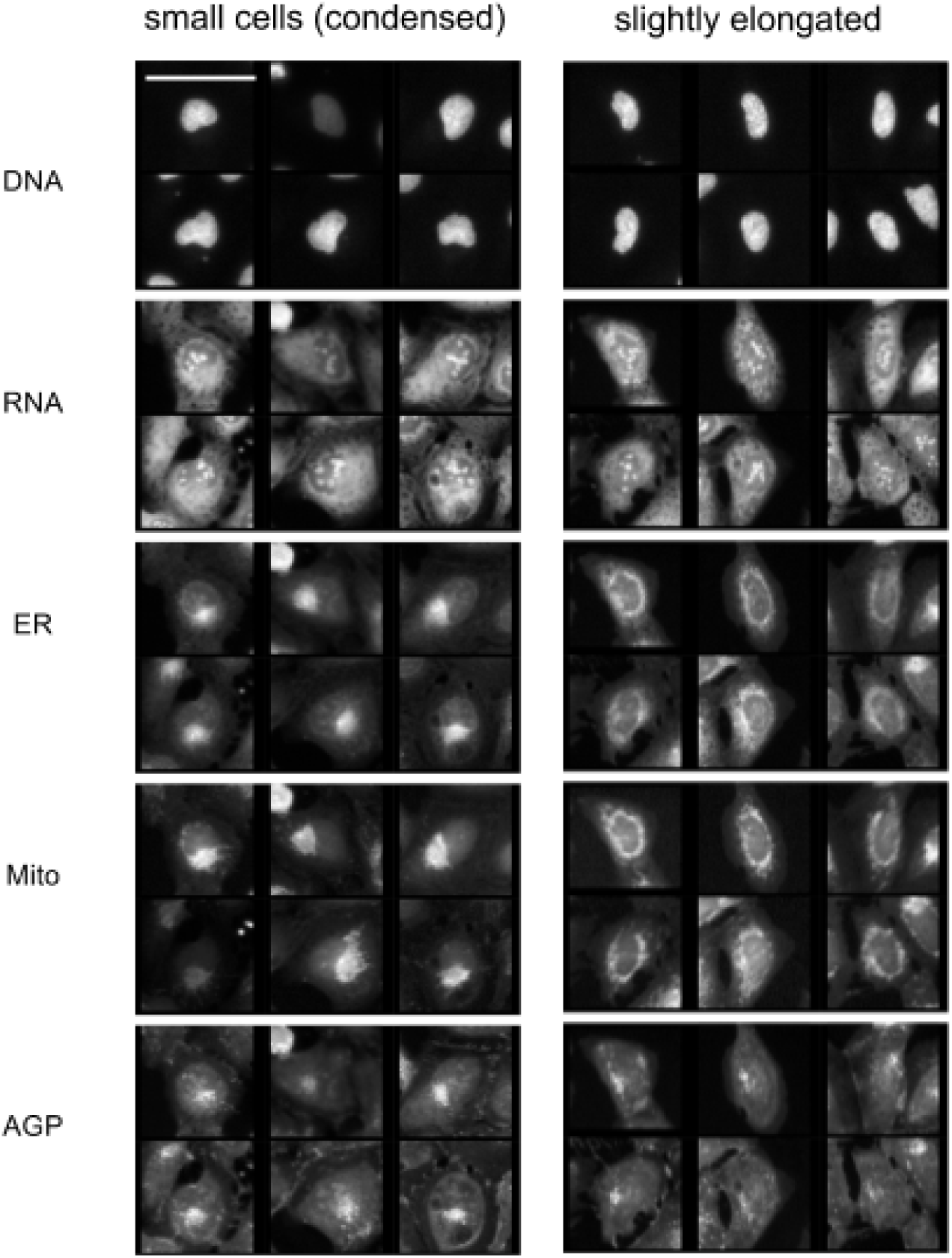
**Common cell subpopulations seen across more than one cluster**. These names are used to annotate clusters of genes in Fig. 3. Example images shown are taken from individual clusters. Scale bar is 63 μ*m* and image intensities are log normalized. References to size and shape in the subpopulation legends refer to both the nucleus and cell borders, unless otherwise noted.

**Supp. Figure 6:**
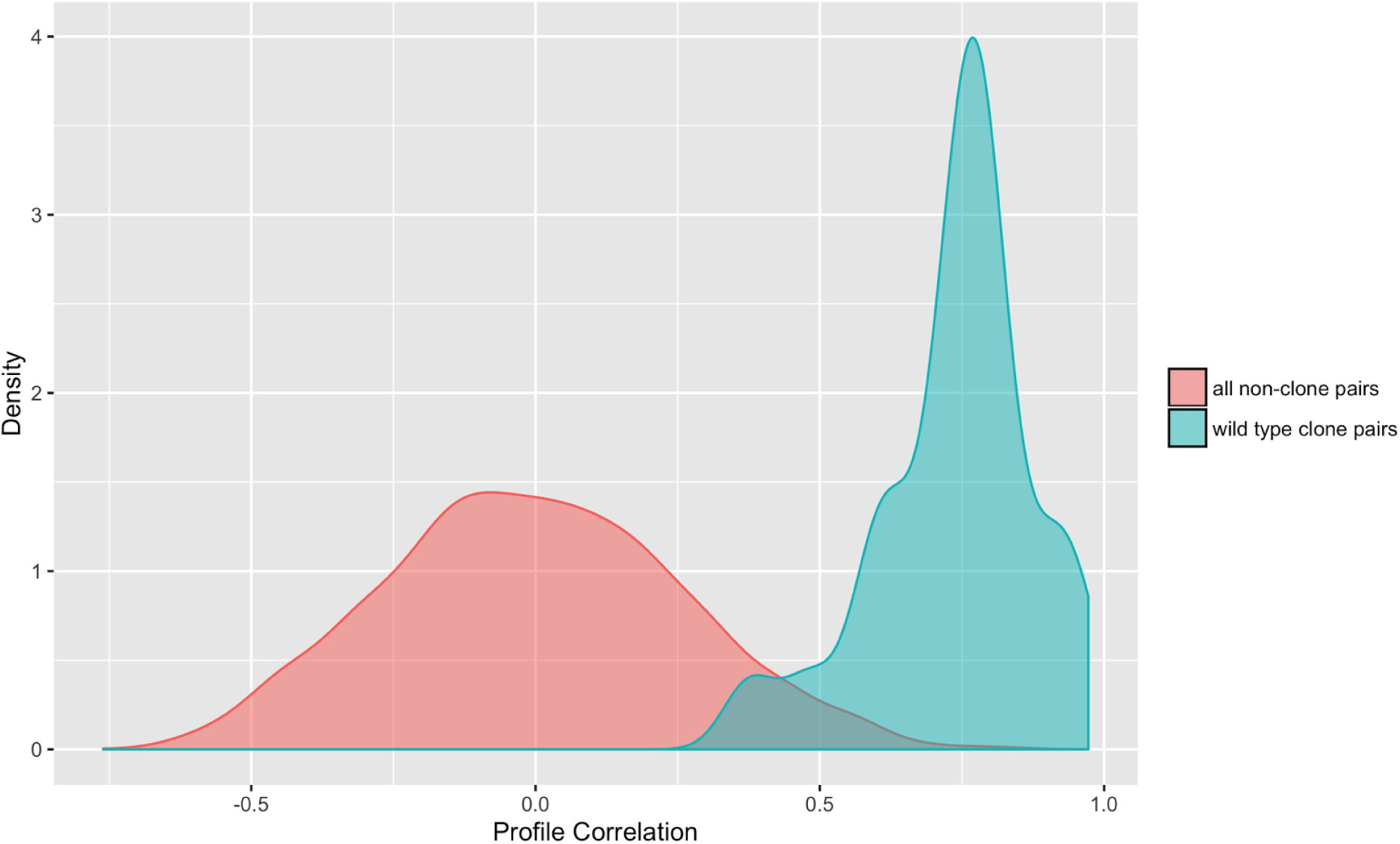
**Selection of the threshold to determine highly correlated gene pairs**. Correlation probability densities of pairs of different genes (labeled ‘non-clone’) and pairs of wild-type clones cross at 0.43, making 0.43 the optimal threshold to determine highly correlated gene pairs.

**Supp. Figure 7:**
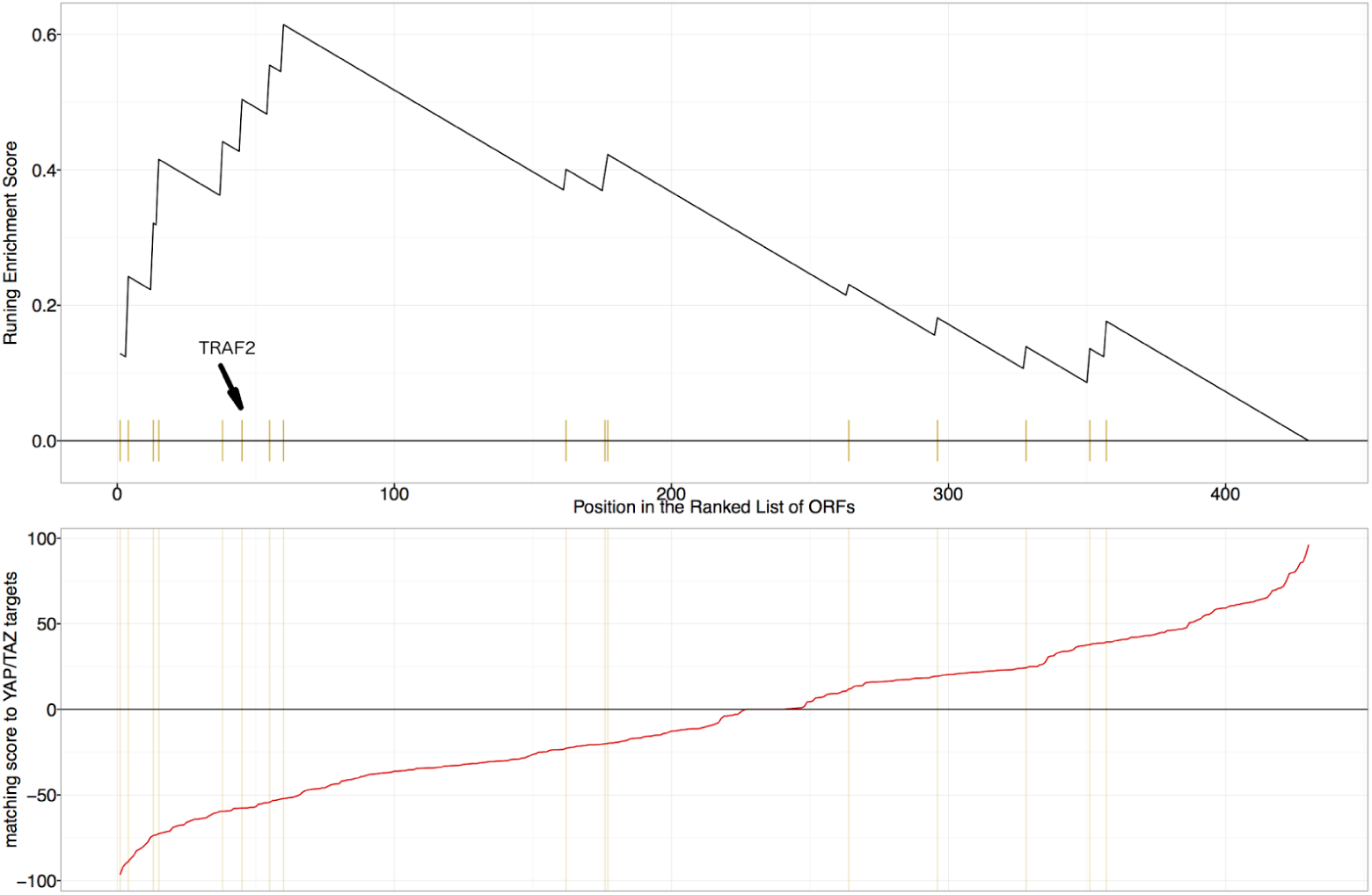
**Gene Set Enrichment Analysis (GSEA) reveals that overexpression constructs sorted based on their similarity to YAP1/WWTR1 overexpression (in terms of impact on particular mRNA targets), are enriched for regulators of the NF-**κ**B pathway (Enrichment Score p-value = 0.0019)**. mRNA targets common to both YAP1 and WWTR overexpression include INPP4B, MAP7, LAMA3, STMN1, and TRAM2, which are positively regulated, and SPP1, IER3, RAB31, and GPR56, which are negatively regulated.

**Supp. Figure 8:**
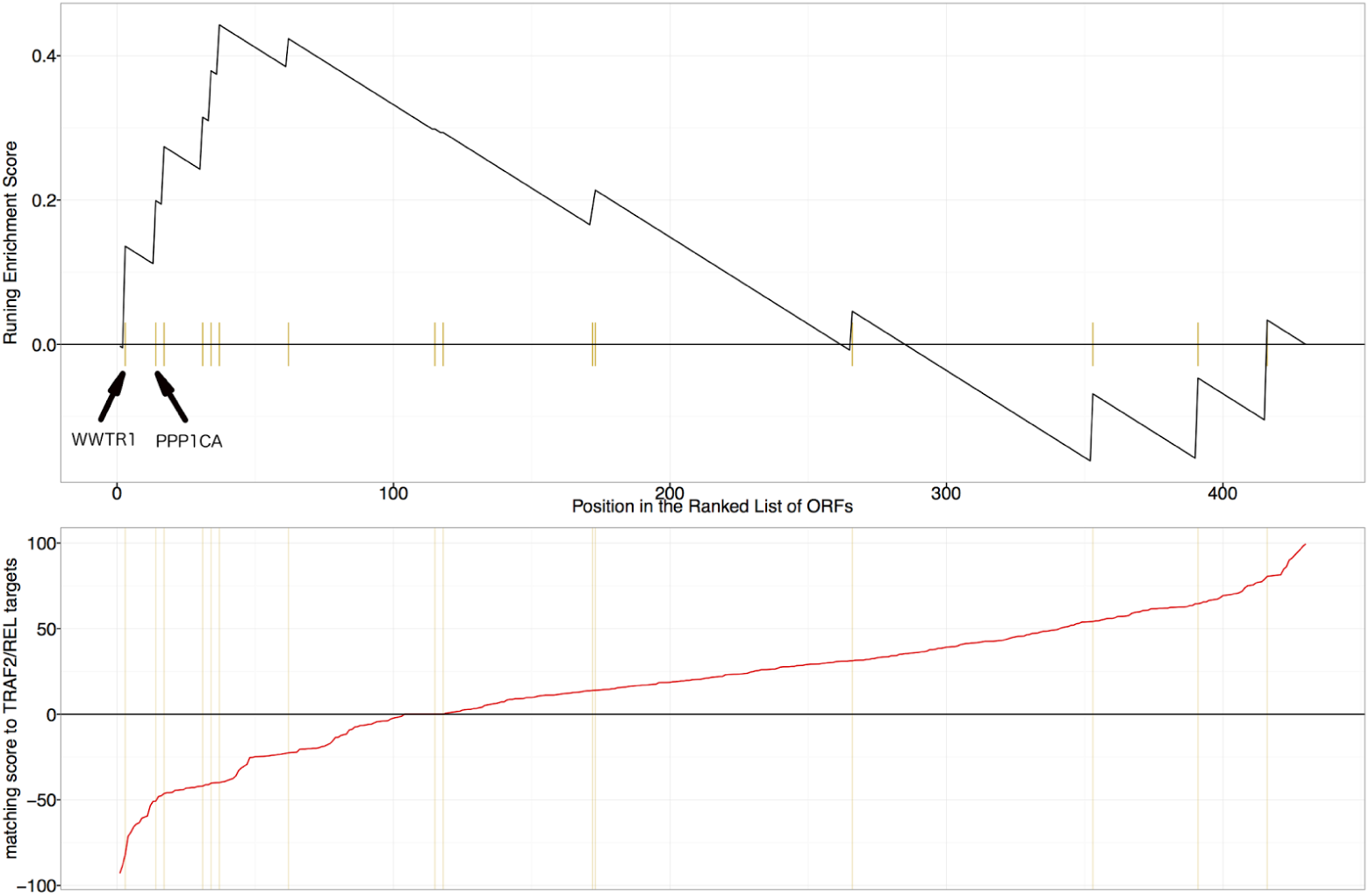
**Gene Set Enrichment Analysis (GSEA) reveals that overexpression constructs sorted based on their similarity to TRAF2/REL overexpression (in terms of impact on particular mRNA targets), are weakly enriched for regulators of the Hippo pathway (Enrichment Score p-value = 0.024)**. mRNA targets common to both TRAF2 and REL overexpression include NFKBIA, IKBKE, AKAP8, and BIRC2, which are positively regulated and RPA3 which is negatively regulated. As compared to Fig. 6C and Supp. Fig. 7, this is a weaker/lower-confidence enrichment - note the lower maximum height (~ 0.44 compared to > 0.6) and higher p-value (0.024 compared to < 0.002). Still, we note that WWTR1 and PPP1CA are the top two matches among those annotated as related to the Hippo pathway in KEGG; PPP1CA (also known as PP-1A) activates TAZ (C.-Y. Liu et al. 2011).

## Supplementary Tables

**Supp. Table 1:**
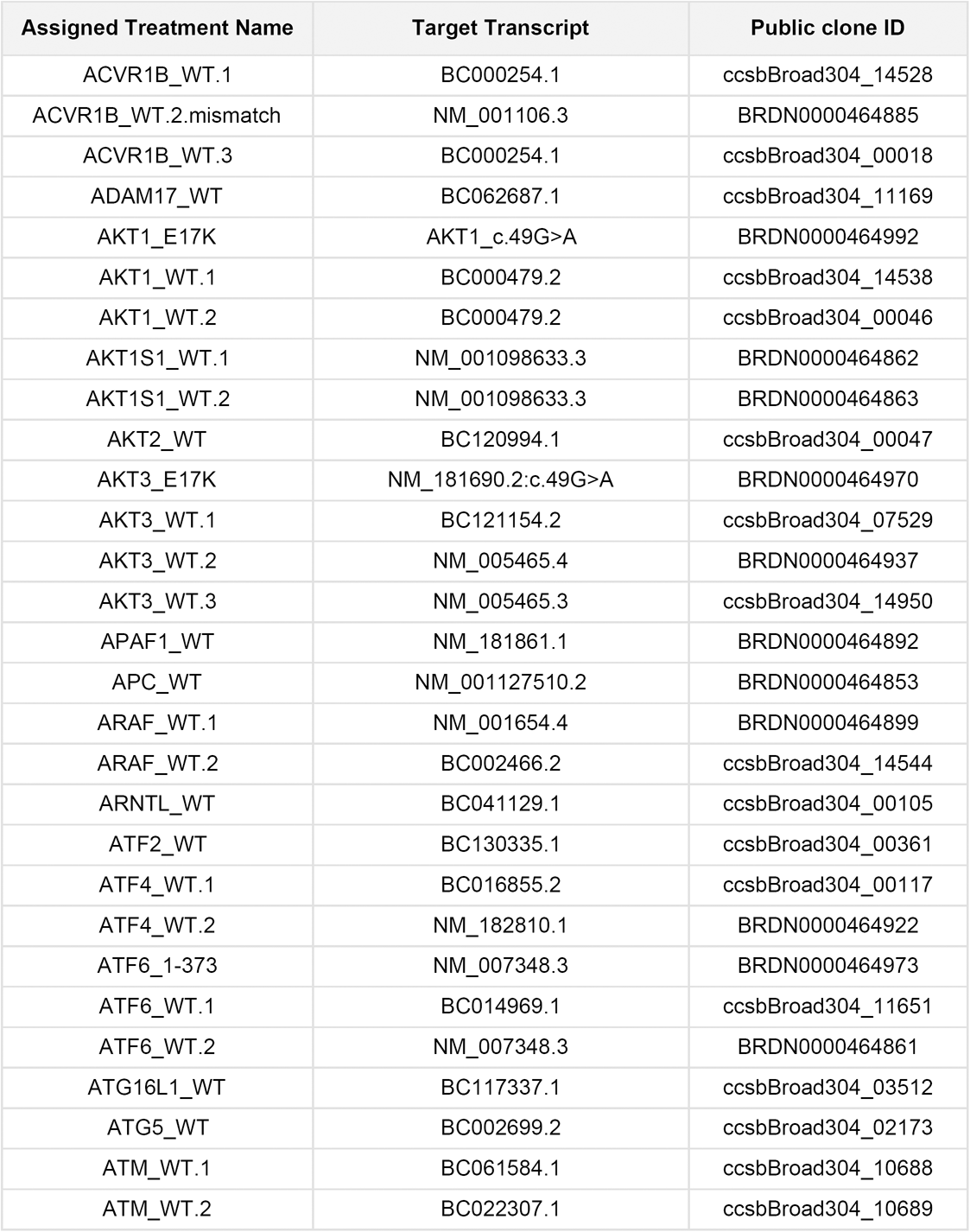
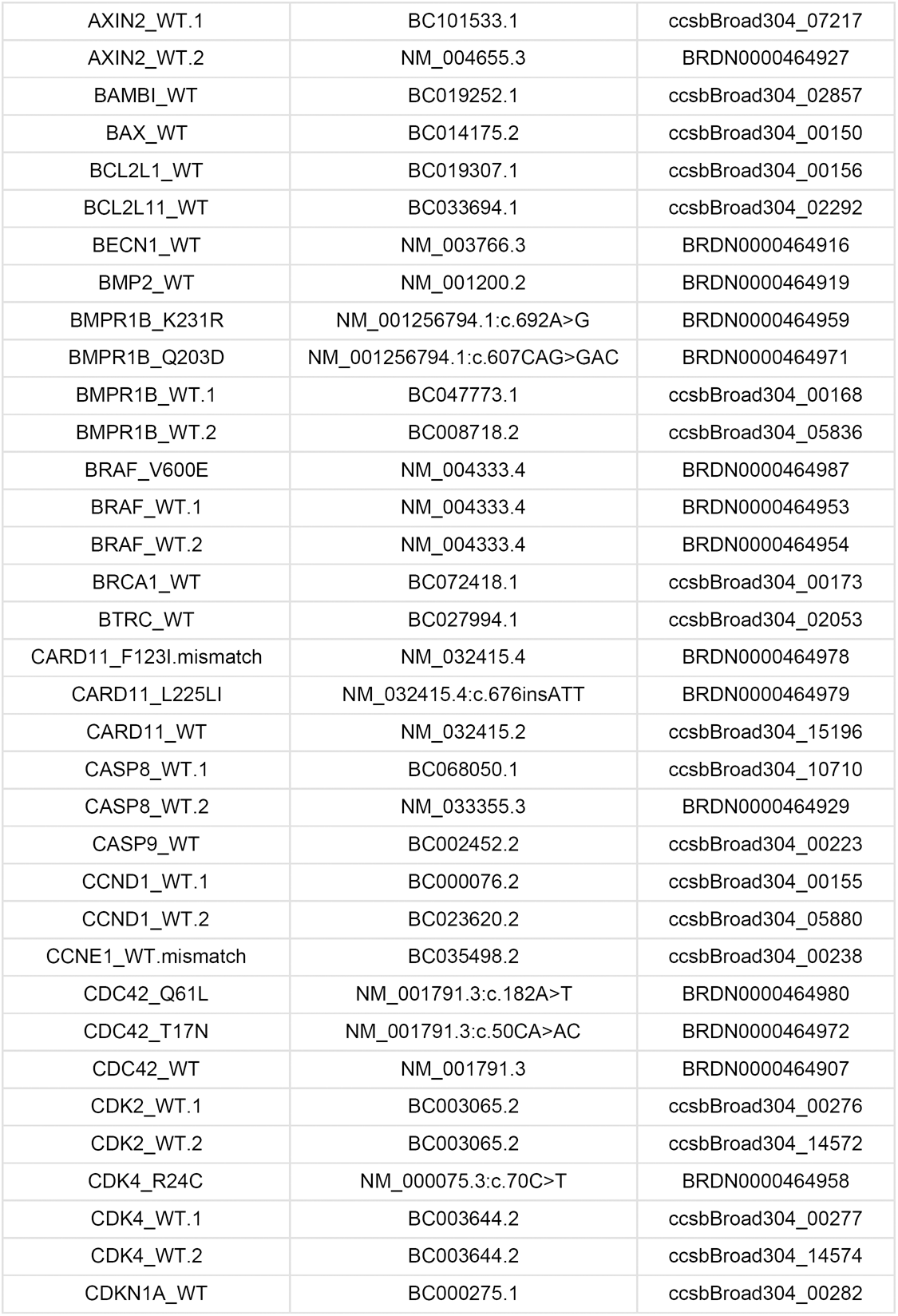
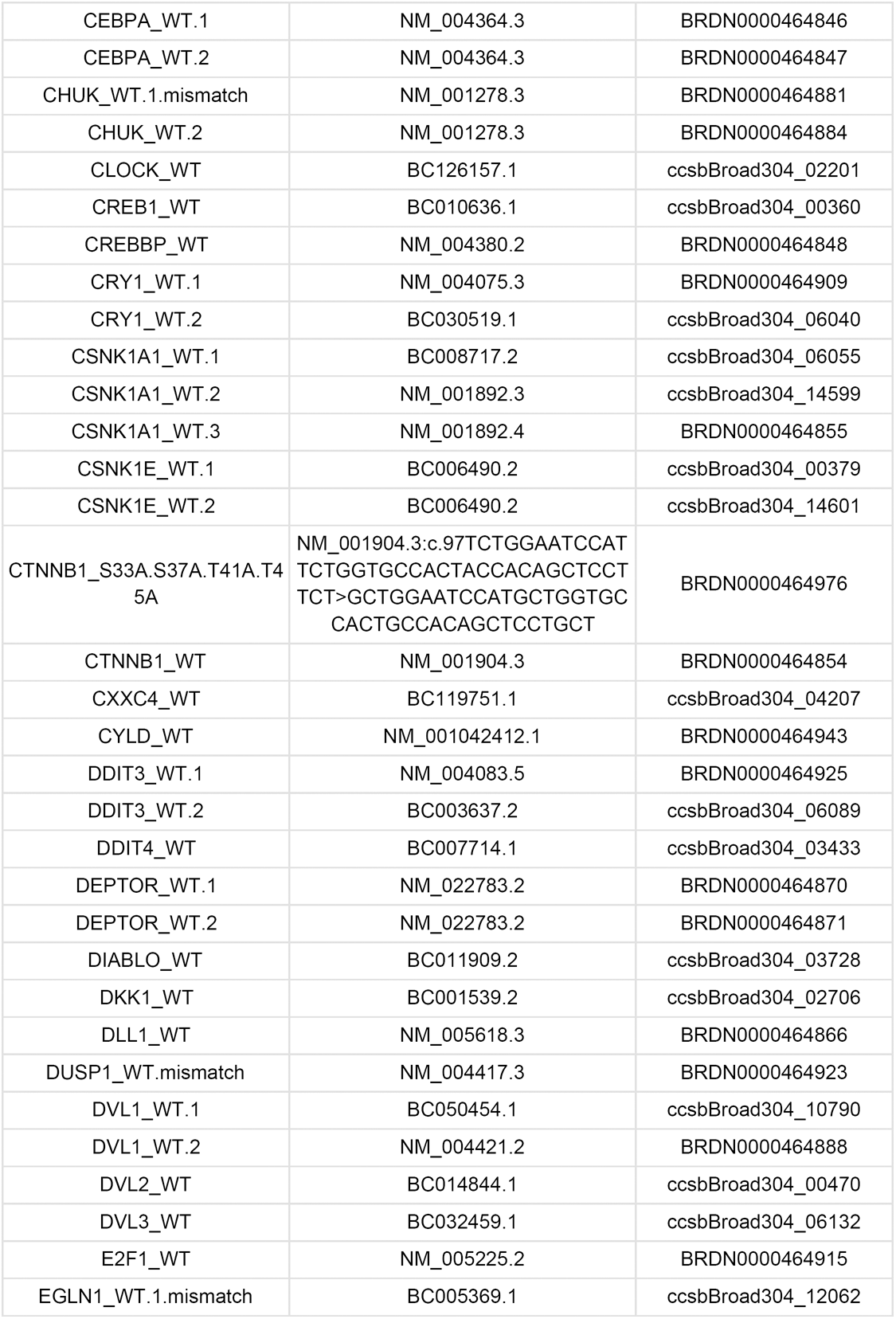
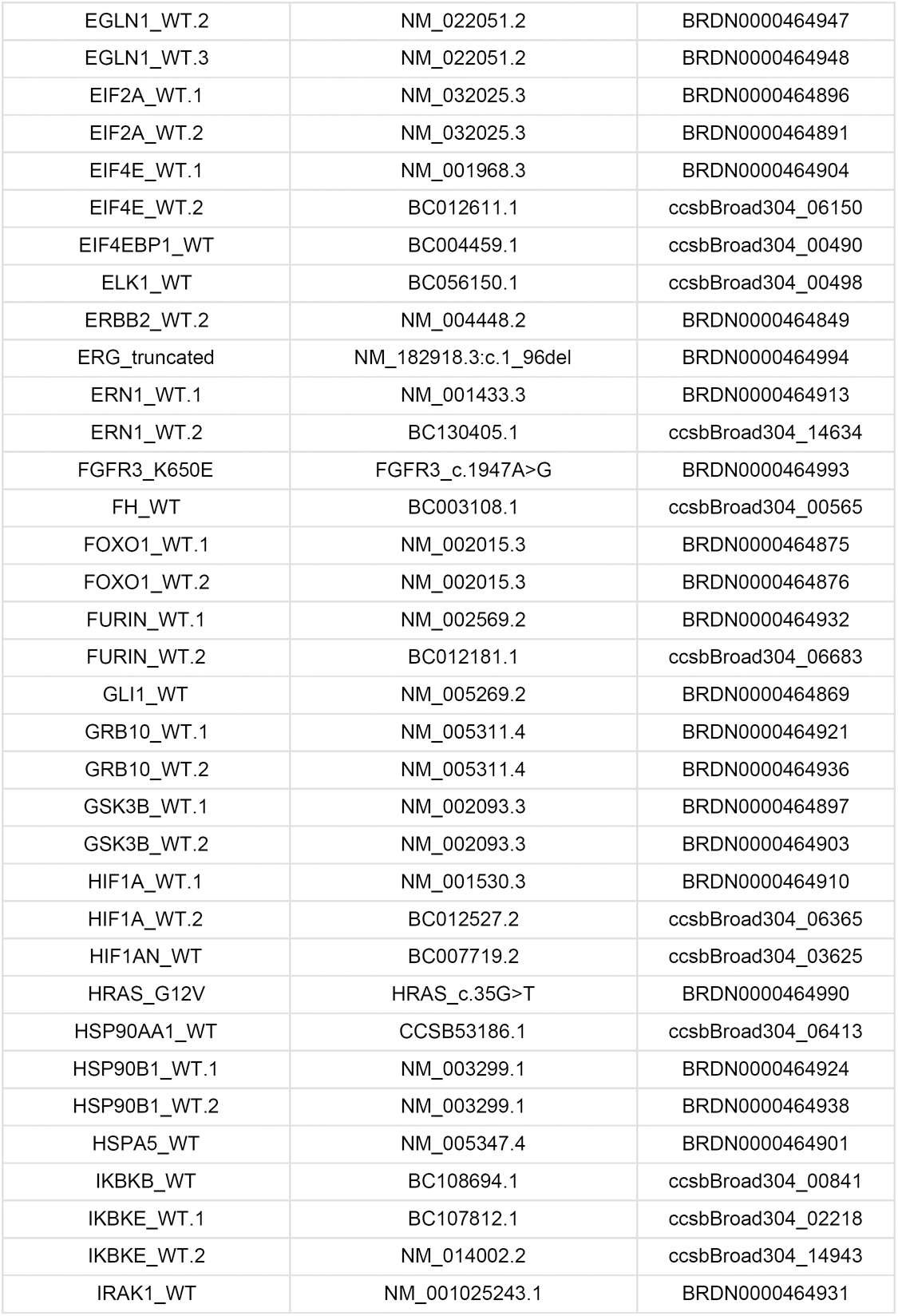
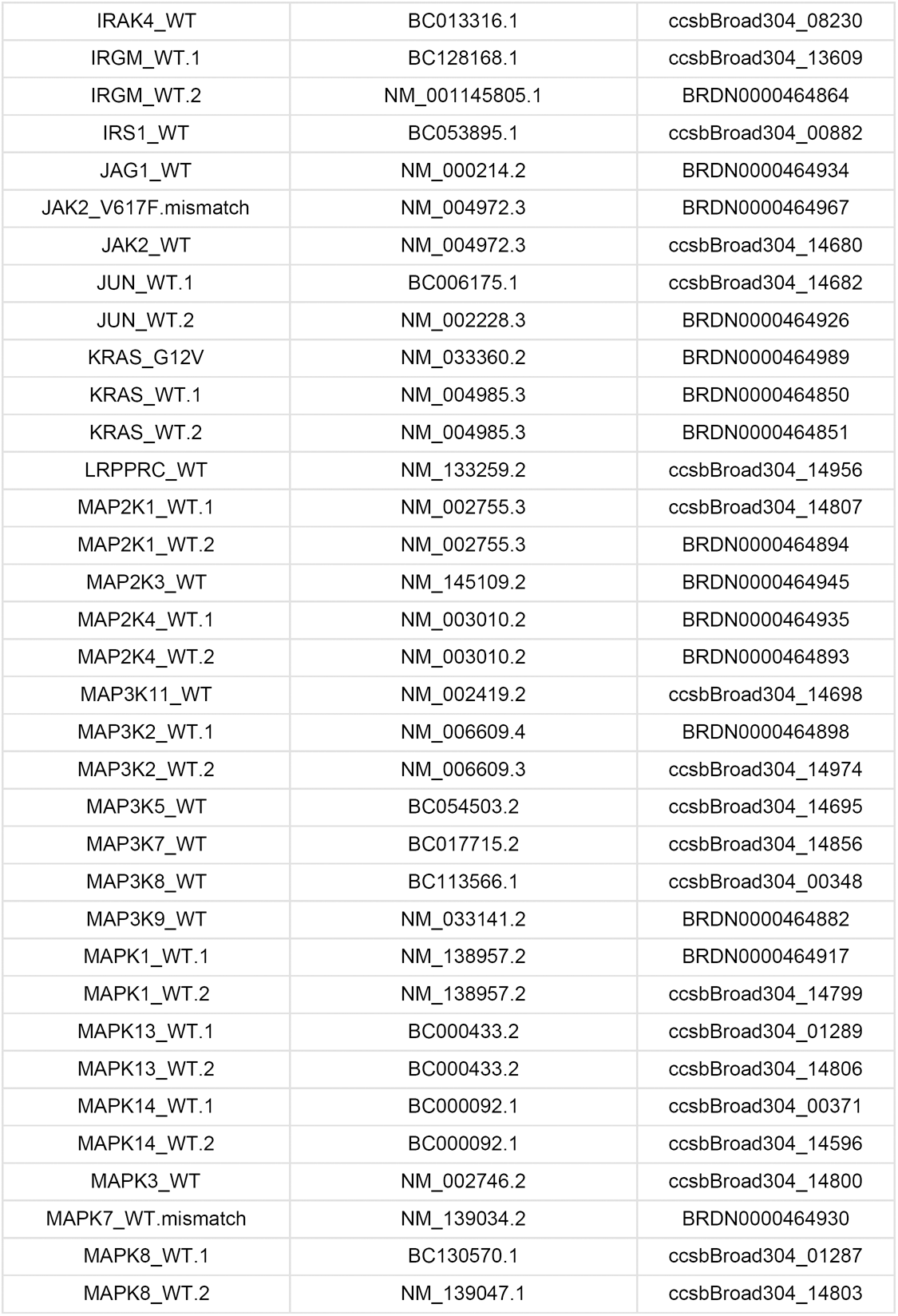
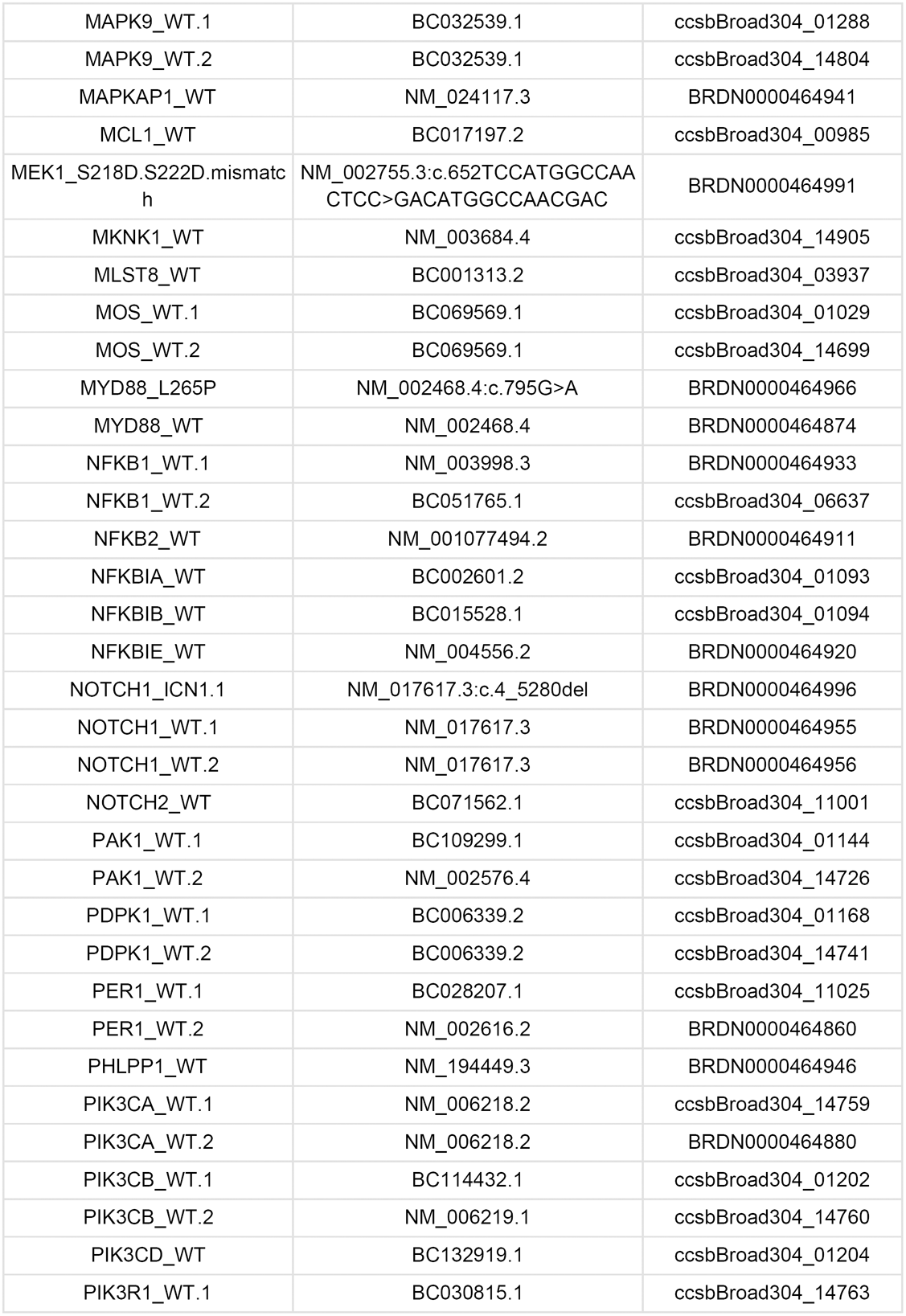
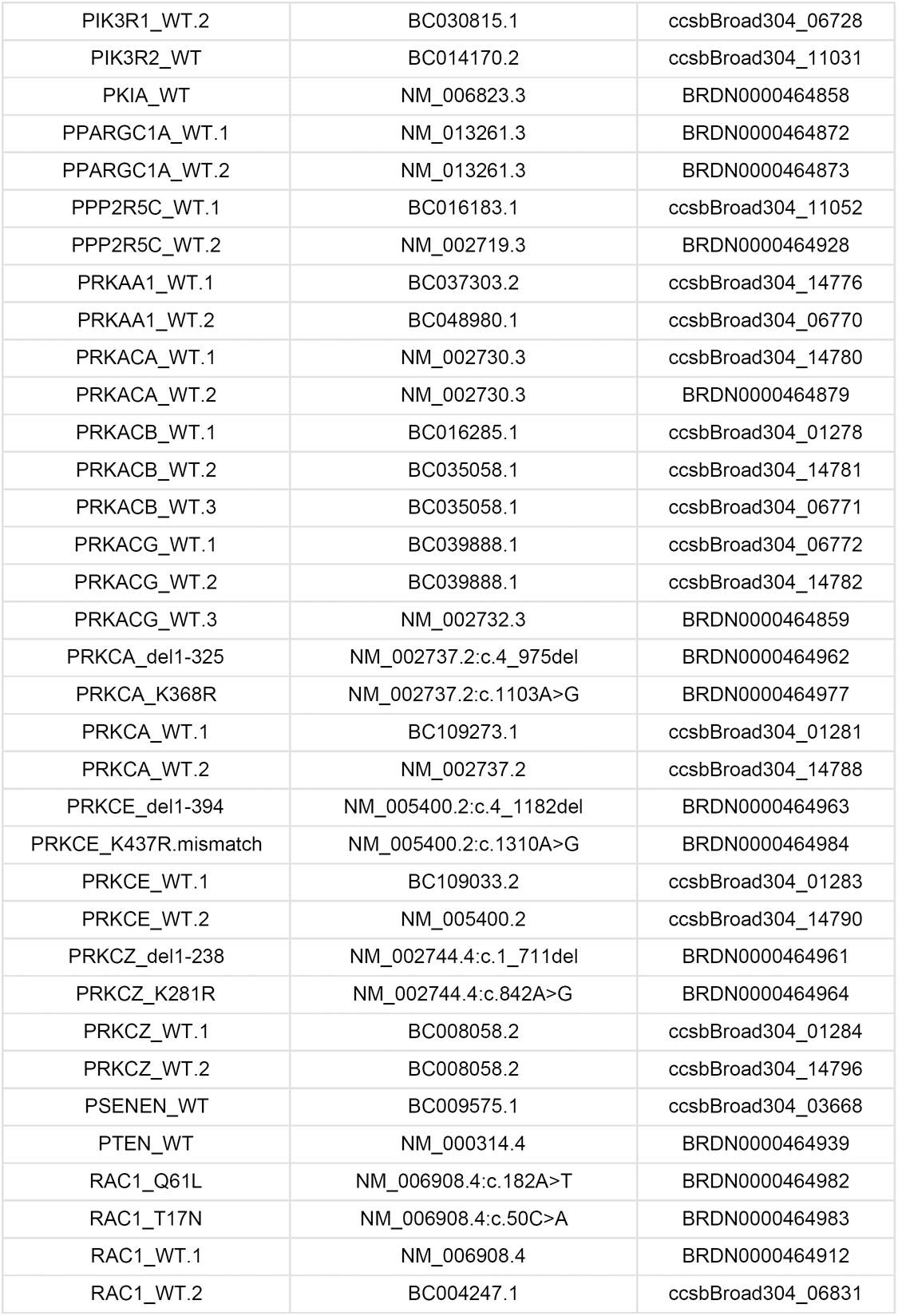
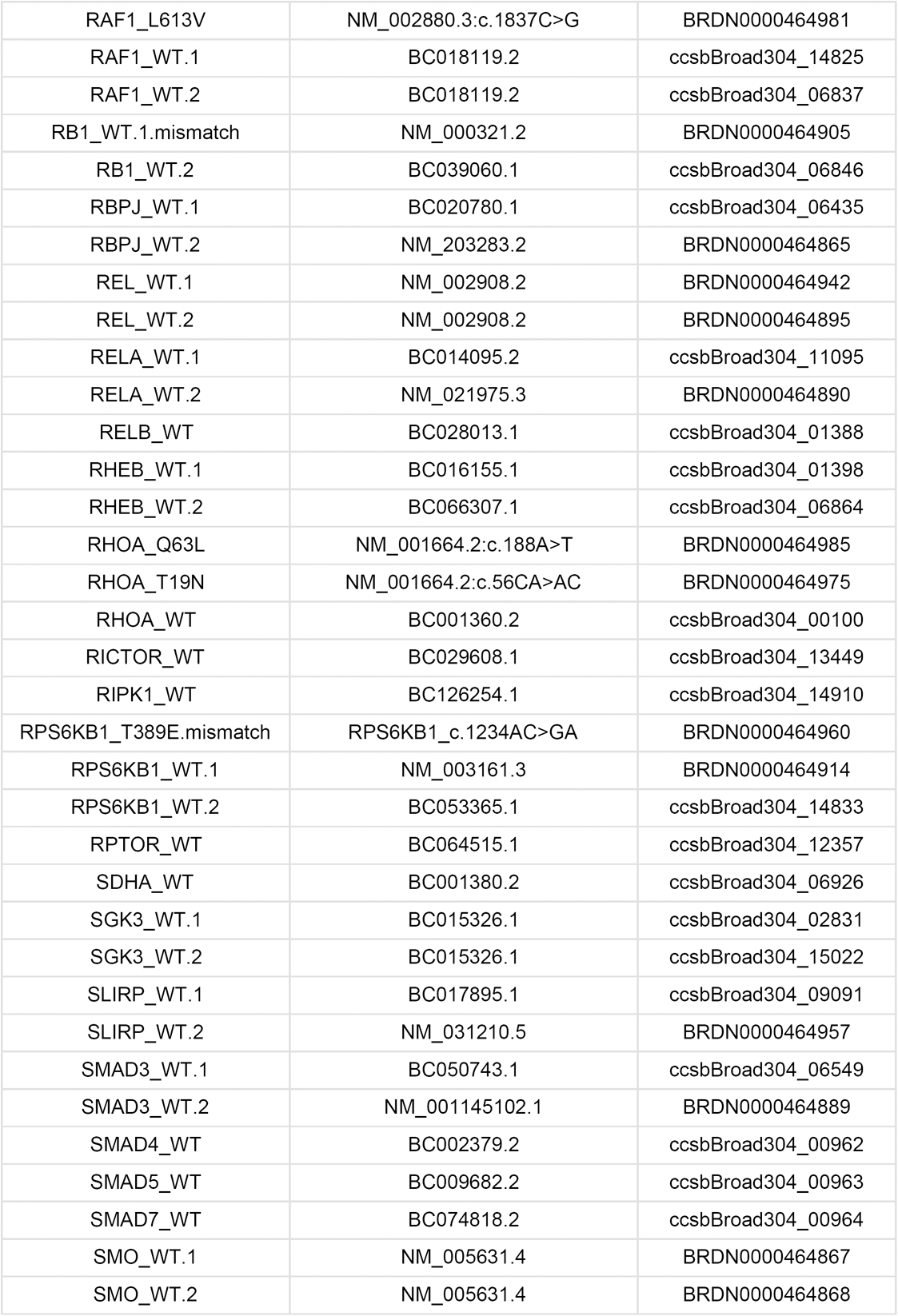
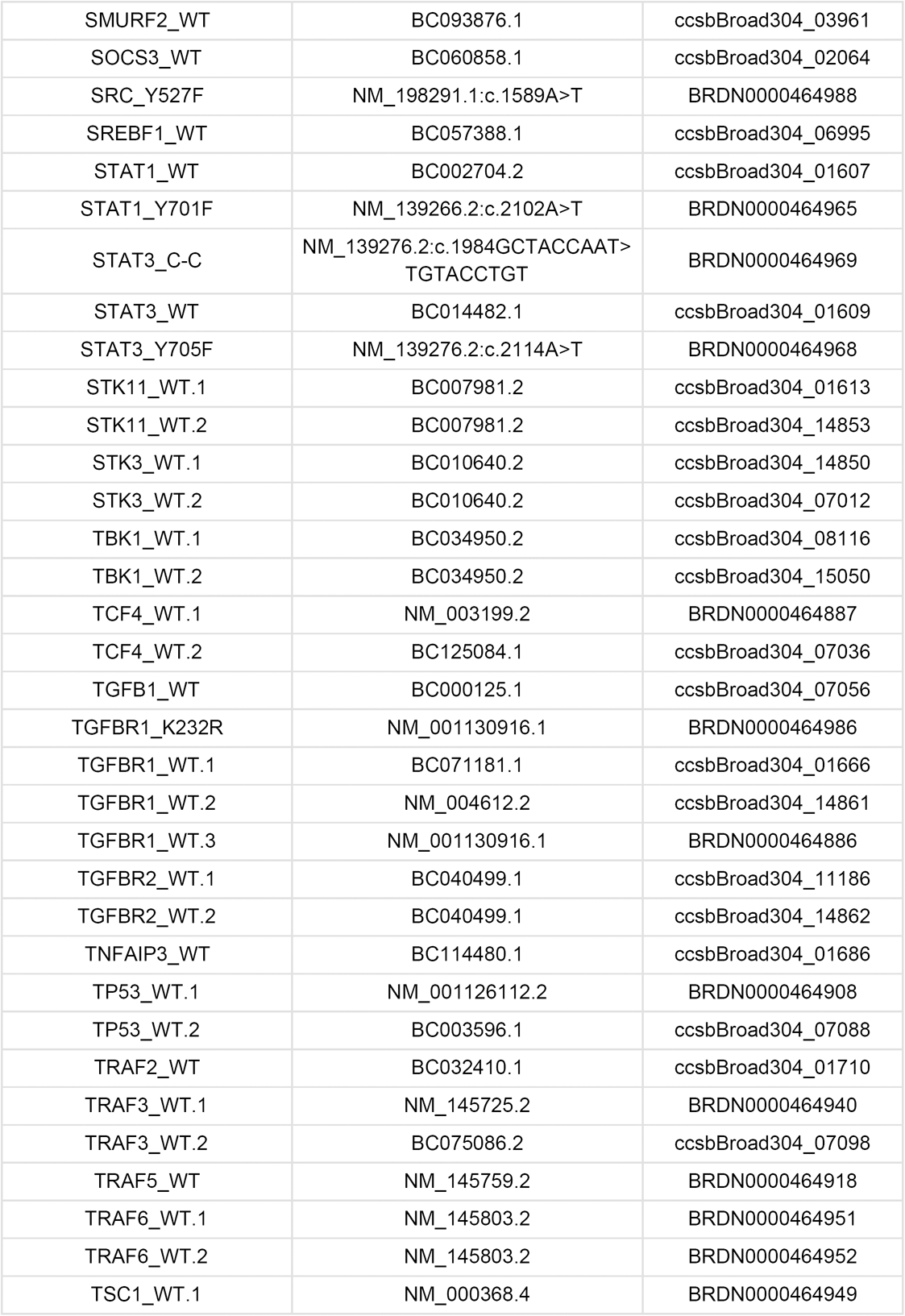
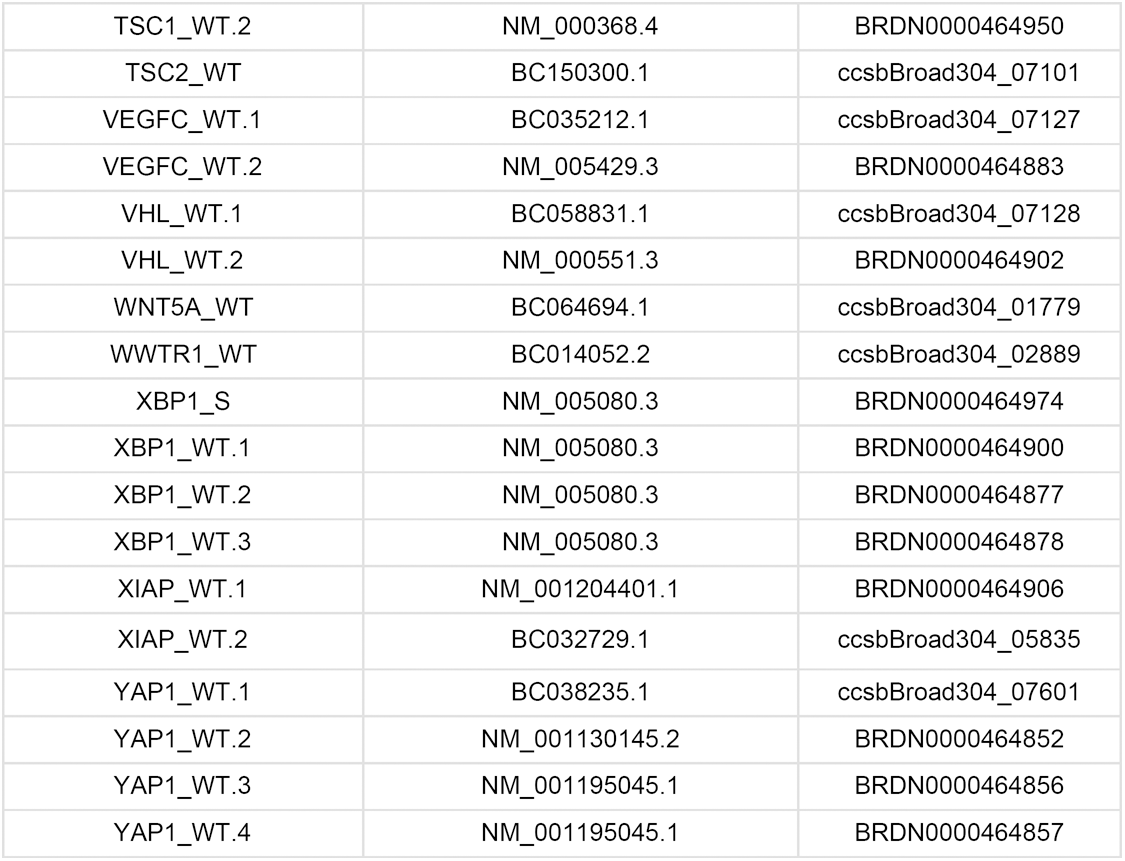
**List of all the 323 constructs used in the experiment along with the target transcript and their public clone ID**.

**Supp. Table 2:** **Replicate correlation is higher in the constitutively active mutant allele compared to the wild-type allele, except for AKT3_E17K. Constitutively active mutant annotations were obtained by literature search for all the mutants in the experiment showing a detectable phenotype. Genes shown here are only those where either the wild-type gene or its constitutively activating allele yielded a phenotype distinct from controls**.

| Gene | Constitutively active mutation | Evidence on being constitutively active | Wild-type median replicate correlation | Mutant median replicate correlation |
| --- | --- | --- | --- | --- |
| BRAF | V600E | (Cantwell-Dorris, O'Leary, and Sheils 2011) | 0.57 | 0.79 |
| RAF1 | L613V | (Wu et al. 2011) | 0.76 | 0.79 |
| KRAS | G12V | (Eser et al. 2014) | 0.43 | 0.72 |
| CDC42 | Q61L | (Nobes and Hall 1999) | 0.64 | 0.89 |
| AKT3 | E17K | (M. S. Kim et al. 2008; M. A. Davies et al. 2008) | 0.84 | 0.70 |
| AKT1 | E17K | (M. S. Kim et al. 2008; M. A. Davies et al. 2008) | 0.40 | 0.93 |
| RHOA | Q63L | (Z.-G. Zhang et al. 2006) | 0.38 | 0.54 |
| RAC1 | Q61L | (Z.-G. Zhang et al. 2006) | 0.30 | 0.53 |

**Supp. Table 3:** **Highly correlated proteins (according to morphology in the Cell Painting assay) that have also been reported to interact physically**. Two genes are highly correlated if their correlation is greater than 0.43 (see Methods).

| Protein 1 | Protein 2 | Method | Evidence |
| --- | --- | --- | --- |
| MAP3K2 | MAP2K3 | Biochemical Activity | Deacon K (1997) |
| HSPA5 | ERN1 | Reconstituted Complex | Liu CY (2003) |
| HSPA5 | ERN1 | Affinity Capture-Western | Yanagisawa K (2010) |
| CDK2 | CCND1 | Affinity Capture-MS | Varjosalo M (2013) |
| CDK2 | CCND1 | Affinity Capture-MS | Hein MY (2015) |
| CCND1 | CDK2 | Affinity Capture-Western | Sweeney KJ (1997) |
| CDK2 | CCND1 | Reconstituted Complex | Higashi H (1996) |
| CDK2 | CCND1 | Affinity Capture-Western | Higashi H (1996) |
| CEBPA | JUN | Affinity Capture-Western | Zada AA (2006) |
| MAP3K7 | TRAF5 | Affinity Capture-Western | Hong S (2007) |
| NFKB1 | RELB | Affinity Capture-MS | Bouwmeester T (2004) |
| RELB | NFKB1 | Affinity Capture-MS | Bouwmeester T (2004) |
| NFKB1 | RELB | Two-hybrid | Wang J (2011) |
| CDC42 | TRAF2 | Affinity Capture-Western | Marivin A (2013) |
| CDC42 | TRAF2 | Reconstituted Complex | Marivin A (2013) |
| KRAS | RAF1 | Two-hybrid | Li W (2000) |
| RAF1 | KRAS | Affinity Capture-Western | Kiyono M (2000) |
| RAF1 | KRAS | Reconstituted Complex | Sui X (2009) |
| RAF1 | KRAS | Affinity Capture-Western | Lito P (2012) |
| KRAS | RAF1 | Affinity Capture-Western | Rajalingam K (2005) |
| KRAS | RAF1 | Affinity Capture-Western | Zhu J (2005) |
| RAF1 | KRAS | Affinity Capture-Western | Blaukat A (2000) |
| RAF1 | KRAS | Affinity Capture-MS | So J (2015) |
| BRAF | RAF1 | Affinity Capture-Western | Weber CK (2001) |
| RAF1 | BRAF | Affinity Capture-Western | Weber CK (2001) |
| RAF1 | BRAF | Affinity Capture-Western | Kilili GK (2010) |
| BRAF | RAF1 | Affinity Capture-Western | Chen C (2008) |
| RAF1 | BRAF | Affinity Capture-Western | Chen C (2008) |
| BRAF | RAF1 | Affinity Capture-Western | Lito P (2012) |
| CSNK1E | STK3 | Affinity Capture-MS | Varjosalo M (2013) |
| RAF1 | STK3 | Affinity Capture-Western | Kilili GK (2010) |
| STK3 | RAF1 | Affinity Capture-Western | Matallanas D (2007) |
| RAF1 | STK3 | Affinity Capture-Western | ONeill E (2004) |
| STK3 | RAF1 | Reconstituted Complex | ONeill E (2004) |
| RAF1 | STK3 | Affinity Capture-MS | ONeill E (2004) |
| STK3 | RAF1 | Affinity Capture-Western | ONeill E (2004) |
| ARAF | HSP90B1 | Affinity Capture-MS | So J (2015) |
| GSK3B | SMURF2 | Affinity Capture-Western | Wu Q (2009) |

**Supp. Table 4:** **Highly correlated genes (according to morphology in the Cell Painting assay) that have also been annotated to be related to the same pathway**. Two genes are highly correlated if their correlation is greater than 0.43 (see Methods).

| Gene 1 | Gene 2 | Pathway annotation |
| --- | --- | --- |
| CCND1_WT.2 | CDK2_WT.2 | Cell Cycle |
| ERN1_WT.1 | HSPA5_WT | ER Stress/UPR |
| BRAF_WT.1 | JUN_WT.1 | MAPK |
| ELK1_WT | JUN_WT.1 | MAPK |
| MAP2K3_WT | MAP2K4_WT.1 | MAPK |
| MAP2K3_WT | MAP3K2_WT.1 | MAPK |
| MAP3K2_WT.1 | MAP3K7_WT | MAPK |
| MAP2K3_WT | MAP3K7_WT | MAPK |
| MAP3K2_WT.1 | MAP3K9_WT | MAPK |
| MAP2K3_WT | MAP3K9_WT | MAPK |
| BRAF_WT.1 | MAP3K9_WT | MAPK |
| ELK1_WT | MAP3K9_WT | MAPK |
| ARAF_WT.1 | MAPK14_WT.1 | MAPK |
| AKT1S1_WT.1 | MLST8_WT | TOR |
| MAP3K9_WT | MOS_WT.1 | MAPK |
| BRAF_WT.1 | MOS_WT.1 | MAPK |
| AKT3_WT.2 | PIK3CB_WT.1 | PI3K/AKT |
| FOXO1_WT.2 | PIK3R1_WT.1 | PI3K/AKT |
| FOXO1_WT.2 | PTEN_WT | PI3K/AKT |
| PIK3R1_WT.1 | PTEN_WT | PI3K/AKT |
| MAP3K9_WT | RAF1_WT.1 | MAPK |
| BRAF_WT.1 | RAF1_WT.1 | MAPK |
| MOS_WT.1 | RAF1_WT.1 | MAPK |
| JUN_WT.1 | RAF1_WT.1 | MAPK |
| NFKB1_WT.1 | RELB_WT | NFkB |
| CDC42_WT | RHOA_WT | Cytoskeletal Re-org |
| NFKB1_WT.1 | TRAF2_WT | NFkB |
| HSP90B1_WT.2 | XBP1_WT.1 | ER Stress/UPR |
| WWTR1_WT | YAP1_WT.1 | Hippo |

**Supp. Table 5:** **Gene Ontology terms associated with each gene cluster**. For clusters containing at least two genes, associated GO terms (limited to those related to Biological Processes) are listed. Mutant and wild-type alleles of a gene are considered the same in the analysis. Clusters containing a single gene are omitted from the table. The p-values are not adjusted and obtained by the Fisher’s exact test, using topGO package in R (Alexa and Rahnenführer 2009).

| Cluster ID | Gene/Alleles | GO Terms |  |  |  |
| --- | --- | --- | --- | --- | --- |
| 2 | TBK1<br>HSP90B1<br>CDK4_R24C<br>STAT3_C-C | GO ID | GO Term | # of genes associated with the GO term in the cluster vs. all the clusters | p-value |
|  |  | GO:0001654 | eye development | 2/3 | 0.014 |
|  |  | GO:0007423 | sensory organ development | 2/7 | 0.086 |
|  |  | GO:0016032 | viral process | 2/7 | 0.086 |
|  |  | GO:0044403 | symbiosis, encompassing mutualism through parasitism | 2/7 | 0.086 |
|  |  | GO:0044419 | interspecies interaction between organisms | 2/7 | 0.086 |
| 3 | CDKN1A<br>KRAS_G12V<br>HRAS_G12V<br>MAP2K4 MAP2K3 | GO ID | GO Term | # of genes associated with the GO term in the cluster vs. all the clusters | p-value |
|  |  | GO:0071900 | regulation of protein serine/threonine kinase activity | 5/17 | 0.0026 |
|  |  | GO:0045860 | positive regulation of protein kinase activity | 5/20 | 0.0066 |
|  |  | GO:0051385 | response to mineralocorticoid | 2/2 | 0.0078 |
|  |  | GO:0043405 | regulation of MAP kinase activity | 4/12 | 0.0086 |
|  |  | GO:0043406 | positive regulation of MAP kinase activity | 4/12 | 0.0086 |
| 4 | MAP3K2<br>CDC42_Q61L<br>MAP3K9 | GO ID | GO Term | # of genes associated with the GO term in the cluster vs. all the clusters | p-value |
|  |  | GO:0032874 | positive regulation of stress-activated MAPK cascade | 3/9 | 0.004 |
|  |  | GO:0046330 | positive regulation of JNK cascade | 3/9 | 0.004 |
|  |  | GO:0070304 | positive regulation of stress-activated protein kinase signaling cascade | 3/9 | 0.004 |
|  |  | GO:0007254 | JNK cascade | 3/10 | 0.0058 |
|  |  | GO:0046328 | regulation of JNK cascade | 3/10 | 0.0058 |
| 5 | XBP1<br>MAPK14<br>RBPJ | GO ID | GO Term | # of genes associated with the GO term in the cluster vs. all the clusters | p-value |
|  |  | GO:0050663 | cytokine secretion | 3/4 | 0.00019 |
|  |  | GO:0051147 | regulation of muscle cell differentiation | 3/4 | 0.00019 |
|  |  | GO:0007517 | muscle organ development | 3/5 | 0.00048 |
|  |  | GO:0051146 | striated muscle cell differentiation | 3/5 | 0.00048 |
|  |  | GO:0051153 | regulation of striated muscle cell differentiation | 3/3 | 4.80E-05 |
| 6 | RAF1_L613V<br>BRAF_V600E | GO ID | GO Term | # of genes associated with the GO term in the cluster vs. all the clusters | p-value |
|  |  | GO:0033135 | regulation of peptidyl-serine phosphorylation | 2/5 | 0.0078 |
|  |  | GO:0033138 | positive regulation of peptidyl-serine phosphorylation | 2/5 | 0.0078 |
|  |  | GO:0023061 | signal release | 2/6 | 0.0118 |
|  |  | GO:0035019 | somatic stem cell population maintenance | 2/6 | 0.0118 |
|  |  | GO:0007411 | axon guidance | 2/8 | 0.022 |
| 7 | RAC1_T17N<br>AKT1_E17K AKT3<br>AKT3_E17K<br>CDC42_T17N | GO ID | GO Term | # of genes associated with the GO term in the cluster vs. all the clusters | p-value |
|  |  | GO:0031294 | lymphocyte costimulation | 3/4 | 0.00076 |
|  |  | GO:0031295 | T cell costimulation | 3/4 | 0.00076 |
|  |  | GO:0045834 | positive regulation of lipid metabolic process | 3/5 | 0.00186 |
|  |  | GO:0010256 | endomembrane system organization | 3/6 | 0.00366 |
|  |  | GO:0030307 | positive regulation of cell growth | 2/2 | 0.00471 |
| 8 | RHOA_Q63L<br>PRKACA<br>GLI1 | GO ID | GO Term | # of genes associated with the GO term in the cluster vs. all the clusters | p-value |
|  |  | GO:0007043 | cell-cell junction assembly | 2/2 | 0.0024 |
|  |  | GO:0043297 | apical junction assembly | 2/2 | 0.0024 |
|  |  | GO:0001503 | ossification | 3/8 | 0.0027 |
|  |  | GO:0007224 | smoothened signaling pathway | 2/3 | 0.007 |
|  |  | GO:0007283 | spermatogenesis | 2/3 | 0.007 |
| 9 | EIF4EBP1 DIABLO | NA |  |  |  |
| 10 | ELK1<br>RHOA | NA |  |  |  |
| 11 | CDC42<br>TRAF2 | GO ID | GO Term | # of genes associated with the GO term in the cluster vs. all the clusters | p-value |
|  |  | GO:0043393 | regulation of protein binding | 2/4 | 0.0047 |
|  |  | GO:0051098 | regulation of binding | 2/7 | 0.0165 |
|  |  | GO:0016567 | protein ubiquitination | 2/8 | 0.022 |
|  |  | GO:0022407 | regulation of cell-cell adhesion | 2/8 | 0.022 |
|  |  | GO:0022409 | positive regulation of cell-cell adhesion | 2/8 | 0.022 |
| 12 | SMAD4 RPS6KB1<br>SMO | GO ID | GO Term | # of genes associated with the GO term in the cluster vs. all the clusters | p-value |
|  |  | GO:0001657 | ureteric bud development | 2/2 | 0.0024 |
|  |  | GO:0001658 | branching involved in ureteric bud morphogenesis | 2/2 | 0.0024 |
|  |  | GO:0001823 | mesonephros development | 2/2 | 0.0024 |
|  |  | GO:0009798 | axis specification | 2/2 | 0.0024 |
|  |  | GO:0009952 | anterior/posterior pattern specification | 2/2 | 0.0024 |
| 13 | CDK2<br>AKT1S1 | NA |  |  |  |
| 14 | PIK3CB<br>TRAF5<br>MAP3K7 | GO ID | GO Term | # of genes associated with the GO term in the cluster vs. all the clusters | p-value |
|  |  | GO:0002250 | adaptive immune response | 2/4 | 0.014 |
|  |  | GO:0050851 | antigen receptor-mediated signaling pathway | 2/5 | 0.023 |
|  |  | GO:0050852 | T cell receptor signaling pathway | 2/5 | 0.023 |
|  |  | GO:0043122 | regulation of I-kappaB kinase/NF-kappaB signaling | 2/6 | 0.033 |
|  |  | GO:0043123 | positive regulation of I-kappaB kinase/NF-kappaB signaling | 2/6 | 0.033 |
| 15 | JUN<br>STK3 | GO ID | GO Term | # of genes associated with the GO term in the cluster vs. all the clusters | p-value |
|  |  | GO:0051098 | regulation of binding | 2/7 | 0.016 |
|  |  | GO:0002520 | immune system development | 2/10 | 0.035 |
|  |  | GO:0008285 | negative regulation of cell proliferation | 2/10 | 0.035 |
|  |  | GO:0030097 | hemopoiesis | 2/10 | 0.035 |
|  |  | GO:0048534 | hematopoietic or lymphoid organ development | 2/10 | 0.035 |
| 16 | PRKCE<br>GRB10 | GO ID | GO Term | # of genes associated with the GO term in the cluster vs. all the clusters | p-value |
|  |  | GO:0010675 | regulation of cellular carbohydrate metabolic process | 2/5 | 0.0078 |
|  |  | GO:0006109 | regulation of carbohydrate metabolic process | 2/6 | 0.0118 |
|  |  | GO:0044262 | cellular carbohydrate metabolic process | 2/6 | 0.0118 |
|  |  | GO:0005996 | monosaccharide metabolic process | 2/7 | 0.0165 |
|  |  | GO:0051051 | negative regulation of transport | 2/8 | 0.022 |
| 18 | CEBPA<br>JUN | GO ID | GO Term | # of genes associated with the GO term in the cluster vs. all the clusters | p-value |
|  |  | GO:0001889 | liver development | 2/3 | 0.0024 |
|  |  | GO:0061008 | hepaticobiliary system development | 2/3 | 0.0024 |
|  |  | GO:0071248 | cellular response to metal ion | 2/3 | 0.0024 |
|  |  | GO:0002573 | myeloid leukocyte differentiation | 2/6 | 0.0118 |
|  |  | GO:0007005 | mitochondrion organization | 2/6 | 0.0118 |
| 19 | KRAS<br>BRAF<br>RAF1<br>MOS | GO ID | GO Term | # of genes associated with the GO term in the cluster vs. all the clusters | p-value |
|  |  | GO:0000186 | activation of MAPKK activity | 4/9 | 0.0005 |
|  |  | GO:0032147 | activation of protein kinase activity | 4/17 | 0.0095 |
|  |  | GO:0007411 | axon guidance | 3/8 | 0.0099 |
|  |  | GO:0097485 | neuron projection guidance | 3/8 | 0.0099 |
|  |  | GO:0007265 | Ras protein signal transduction | 3/9 | 0.0146 |
| 20 | YAP1<br>WWTR1 | GO ID | GO Term | # of genes associated with the GO term in the cluster vs. all the clusters | p-value |
|  |  | GO:0035850 | epithelial cell differentiation involved in kidney development | 2/2 | 0.00078 |
|  |  | GO:0061005 | cell differentiation involved in kidney development | 2/2 | 0.00078 |
|  |  | GO:0072160 | nephron tubule epithelial cell differentiation | 2/2 | 0.00078 |
|  |  | GO:0072170 | metanephric tubule development | 2/2 | 0.00078 |
|  |  | GO:0072182 | regulation of nephron tubule epithelial cell differentiation | 2/2 | 0.00078 |
| 21 | PIK3R1<br>PTEN |  |  |  |  |
|  |  | GO ID | GO Term | # of genes associated with the GO term in the cluster vs. all the clusters | p-value |
|  |  | GO:0001953 | negative regulation of cell-matrix adhesion | 2/2 | 0.00078 |
|  |  | GO:0007162 | negative regulation of cell adhesion | 2/2 | 0.00078 |
|  |  | GO:0010812 | negative regulation of cell-substrate adhesion | 2/2 | 0.00078 |
|  |  | GO:0006650 | glycerophospholipid metabolic process | 2/3 | 0.00235 |
|  |  | GO:0006661 | phosphatidylinositol biosynthetic process | 2/3 | 0.00235 |
| 23 | NFKB1<br>PRKACG |  |  |  |  |
|  |  | GO ID | GO Term | # of genes associated with the GO term in the cluster vs. all the clusters | p-value |
|  |  | GO:0044283 | small molecule biosynthetic process | 2/5 | 0.0078 |
|  |  | GO:0016051 | carbohydrate biosynthetic process | 2/6 | 0.0118 |
|  |  | GO:0002220 | innate immune response activating cell surface receptor signaling pathway | 2/7 | 0.0165 |
|  |  | GO:0002223 | stimulatory C-type lectin receptor signaling pathway | 2/7 | 0.0165 |
|  |  | GO:0005975 | carbohydrate metabolic process | 2/10 | 0.0353 |
| 24 | SMURF2<br>DVL3 | GO ID | GO Term | # of genes associated with the GO term in the cluster vs. all the clusters | p-value |
|  |  | GO:0030177 | positive regulation of Wnt signaling pathway | 2/5 | 0.0078 |
|  |  | GO:0090263 | positive regulation of canonical Wnt signaling pathway | 2/5 | 0.0078 |
|  |  | GO:0060828 | regulation of canonical Wnt signaling pathway | 2/10 | 0.0353 |
|  |  | GO:0030111 | regulation of Wnt signaling pathway | 2/11 | 0.0431 |
|  |  | GO:0060070 | canonical Wnt signaling pathway | 2/12 | 0.0518 |
| 25 | PRKCZ_del1-238<br>FOXO1 | GO ID | GO Term | # of genes associated with the GO term in the cluster vs. all the clusters | p-value |
|  |  | GO:0001933 | negative regulation of protein phosphorylation | 2/9 | 0.028 |
|  |  | GO:0031400 | negative regulation of protein modification process | 2/9 | 0.028 |
|  |  | GO:0010563 | negative regulation of phosphorus metabolic process | 2/11 | 0.043 |
|  |  | GO:0042326 | negative regulation of phosphorylation | 2/11 | 0.043 |
|  |  | GO:0045936 | negative regulation of phosphate metabolic process | 2/11 | 0.043 |

**Supp. Table 6:** **Rank ordered list of distinctive features based on their z-scores for Cluster** 19. The top 50 features with highest absolute z-score are shown.

| Feature name | z-score |
| --- | --- |
| mad_Cells_Intensity_LowerQuartileIntensity_RNA | 8.91 |
| mad_Cytoplasm_Intensity_LowerQuartileIntensity_RNA | 8.6 |
| mad_Cytoplasm_Intensity_MeanIntensity_RNA | 8.53 |
| mad_Cytoplasm_Intensity_MedianIntensity_RNA | 8.45 |
| Cytoplasm_Intensity_LowerQuartileIntensity_RNA | 8.34 |
| Cells_Intensity_LowerQuartileIntensity_RNA | 8.17 |
| Cells_Intensity_MeanIntensityEdge_RNA | 7.74 |
| mad_Cells_Intensity_MeanIntensity_RNA | 7.61 |
| mad_Cells_Intensity_MedianIntensity_RNA | 7.31 |
| mad_Cytoplasm_Intensity_UpperQuartileIntensity_AGP | 7.26 |
| Cells_Intensity_StdIntensityEdge_RNA | 7.19 |
| mad_Cytoplasm_Intensity_MeanIntensity_AGP | 7.05 |
| mad_Cells_Intensity_MedianIntensity_AGP | 7 |
| mad_Cells_Intensity_MeanIntensityEdge_RNA | 6.74 |
| mad_Cytoplasm_Intensity_MedianIntensity_AGP | 6.73 |
| mad_Cells_Intensity_UpperQuartileIntensity_RNA | 6.63 |
| mad_Cytoplasm_Intensity_UpperQuartileIntensity_RNA | 6.56 |
| mad_Cytoplasm_Intensity_MeanIntensityEdge_RNA | 6.44 |
| Cytoplasm_Intensity_MeanIntensityEdge_RNA | 6.36 |
| Cytoplasm_Intensity_MedianIntensity_RNA | 6.27 |
| Cells_Intensity_MaxIntensityEdge_RNA | 6.25 |
| mad_Cells_Intensity_MeanIntensity_AGP | 6.17 |
| mad_Cells_Intensity_UpperQuartileIntensity_AGP | 6.12 |
| mad_Cytoplasm_Intensity_MeanIntensity_Mito | 6.09 |
| mad_Cytoplasm_Intensity_LowerQuartileIntensity_Mito | 5.98 |
| mad_Cells_Intensity_LowerQuartileIntensity_Mito | 5.89 |
| mad_Cells_Intensity_MeanIntensity_Mito | 5.72 |
| mad_Cells_Intensity_StdIntensityEdge_RNA | 5.69 |
| mad_Cytoplasm_Intensity_MedianIntensity_Mito | 5.53 |
| mad_Cytoplasm_RadialDistribution_MeanFrac_Mito_1of4 | 5.53 |
| mad_Cytoplasm_RadialDistribution_MeanFrac_Mito_4of4 | 5.52 |
| mad_Cytoplasm_Intensity_StdIntensity_Mito | 5.5 |
| Cells_Intensity_MedianIntensity_RNA | 5.48 |
| Cytoplasm_Intensity_MeanIntensity_RNA | 5.47 |
| mad_Cytoplasm_RadialDistribution_MeanFrac_Mito_2of4 | 5.46 |
| mad_Nuclei_Intensity_MeanIntensityEdge_RNA | 5.37 |
| Cytoplasm_RadialDistribution_MeanFrac_Mito_4of4 | -5.36 |
| mad_Nuclei_Intensity_MedianIntensity_RNA | 5.32 |
| Cells_Intensity_MeanIntensity_RNA | 5.23 |
| mad_Cytoplasm_Intensity_MeanIntensityEdge_AGP | 5.21 |
| mad_Cells_Intensity_LowerQuartileIntensity_ER | 5.17 |
| mad_Nuclei_Intensity_MeanIntensity_RNA | 5.15 |
| Cytoplasm_RadialDistribution_FracAtD_Mito_2of4 | 5.15 |
| mad_Cytoplasm_Intensity_StdIntensityEdge_RNA | 5.11 |
| mad_Cells_Intensity_MedianIntensity_Mito | 5.04 |
| mad_Cells_Intensity_MeanIntensityEdge_Mito | 5.04 |
| mad_Cells_Intensity_StdIntensityEdge_AGP | 5.02 |
| mad_Cells_Intensity_StdIntensityEdge_Mito | 4.99 |
| mad_Cytoplasm_Intensity_MedianIntensity_ER | 4.95 |
| Cytoplasm_RadialDistribution_MeanFrac_AGP_4of4 | -4.93 |

**Supp. Table 7:**
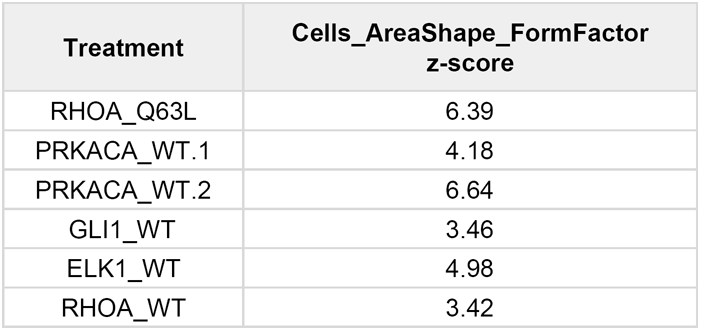
**All genes/alleles in Cluster 8 and 10 induce cell rounding**. “Form Factor” is a measurement in the morphological profile that is defined as the area of an object divided by its perimeter; the metric shown here was calculated on the shape of the entire cell, which is defined by RNA staining (SYT014). The positive z-scores seen here indicate increased cell rounding.

| Treatment | Cells_AreaShape_FormFactor<br>z-score |
| --- | --- |
| RHOA_Q63L | 6.39 |
| PRKACA_WT.1 | 4.18 |
| PRKACA_WT.2 | 6.64 |
| GLI1_WT | 3.46 |
| ELK1_WT | 4.98 |
| RHOA_WT | 3.42 |

**Supp. Table 8:** **The NF-**κ**B signaling pathway is the most enriched when searching for gene overexpressions that downregulate known YAP/TAZ targets (CYR61, CTGF, and BIRC5)**. The ORFs corresponding to each pathway are listed in the last column.

| Pathway Name | Adjusted p-value | Gene IDs |
| --- | --- | --- |
| NF-kappa B signaling pathway | 2e-08 | BCL10/GADD45B/CD40/LTBR/TIRAP/RELB/TRAF2/LYN |
| Epstein-Barr virus infection | 5.3e-06 | CDKN1B/IFNG/CDKN1A/CD40/RELB/MAP2K6/TRAF2/LYN |
| Cell cycle | 0.0011 | CDKN2C/CDKN1B/CDKN1A/GADD45B/GADD45A |
| Osteoclast differentiation | 0.0011 | IFNB1/IFNG/RELB/MAP2K6/TRAF2 |
| FoxO signaling pathway | 0.0011 | CDKN1B/CDKN1A/GADD45B/RAF1/GADD45A |
| Hepatitis B | 0.0013 | CDKN1B/IFNB1/CDKN1A/TIRAP/RAF1 |
| MAPK signaling pathway | 0.0013 | GADD45B/RELB/RAF1/MAP2K6/TRAF2/GADD45A |
| HTLV-I infection | 0.0013 | CDKN2C/CDKN1A/CD40/LTBR/RELB/IL2RB |
| Jak-STAT signaling pathway | 0.0013 | IFNB1/IFNG/CDKN1A/RAF1/IL2RB |
| Tuberculosis | 0.002 | IFNB1/IFNG/BCL10/TIRAP/RAF1 |
| Transcriptional misregulation in cancer | 0.002 | CDKN2C/CDKN1B/CDKN1A/CD40/IL2RB |
| Endocrine resistance | 0.002 | CDKN2C/CDKN1B/CDKN1A/RAF1 |
| HIF-1 signaling pathway | 0.0023 | CDKN1B/IFNG/CDKN1A/LTBR |
| Toll-like receptor signaling pathway | 0.0023 | IFNB1/CD40/TIRAP/MAP2K6 |
| Viral carcinogenesis | 0.0026 | CDKN1B/CDKN1A/LTBR/TRAF2/LYN |
| Hepatitis C | 0.0049 | IFNB1/CDKN1A/RAF1/TRAF2 |
| Measles | 0.005 | CDKN1B/IFNB1/IFNG/IL2RB |
| Apoptosis | 0.0053 | GADD45B/RAF1/TRAF2/GADD45A |
| Cytokine-cytokine receptor interaction | 0.0067 | IFNB1/IFNG/CD40/LTBR/IL2RB |
| p53 signaling pathway | 0.0069 | CDKN1A/GADD45B/GADD45A |
| Fc epsilon RI signaling pathway | 0.0069 | RAF1/MAP2K6/LYN |
| B cell receptor signaling pathway | 0.0072 | BCL10/RAF1/LYN |
| Chronic myeloid leukemia | 0.0072 | CDKN1B/CDKN1A/RAF1 |
| Influenza A | 0.009 | IFNB1/IFNG/RAF1/MAP2K6 |
| ErbB signaling pathway | 0.011 | CDKN1B/CDKN1A/RAF1 |
| Prostate cancer | 0.011 | CDKN1B/CDKN1A/RAF1 |
| PI3K-Akt signaling pathway | 0.014 | CDKN1B/IFNB1/CDKN1A/RAF1/IL2RB |
| T cell receptor signaling pathway | 0.017 | IFNG/BCL10/RAF1 |
| Toxoplasmosis | 0.022 | IFNG/CD40/MAP2K6 |
| Allograft rejection | 0.024 | IFNG/CD40 |
| Bladder cancer | 0.027 | CDKN1A/RAF1 |
| Natural killer cell mediated cytotoxicity | 0.029 | IFNB1/IFNG/RAF1 |
| Breast cancer | 0.033 | CDKN1A/RAF1/HEY1 |
| Intestinal immune network for IgA production | 0.033 | CD40/LTBR |
| Malaria | 0.033 | IFNG/CD40 |
| Long-term depression | 0.048 | RAF1/LYN |

## References

Abell Kathrine, Antonio Bilancio, Richard W. E. Clarkson, Paul G. Tiffen, Anton I. Altaparmakov, Thomas G. Burdon, Tomoichiro Asano, Bart Vanhaesebroeck, and Christine J. Watson. 2005. “Stat3-Induced Apoptosis Requires a Molecular Switch in PI (3) K Subunit Composition.” Nature Cell Biology 7 (4). Nature Publishing Group: 392-98. http://www.nature.com/ncb/journal/v7/n4/abs/ncb1242.html.

Alexa Adrian, and Jörg Rahnenführer. 2009. “Gene Set Enrichment Analysis with topGO.” Available. https://bioconductor.riken.jp/packages/3.2/bioc/vignettes/topGO/inst/doc/topGO.pdf.

Amberger Joanna S., Carol A. Bocchini, François Schiettecatte, Alan F. Scott, and Ada Hamosh. 2015. “OMIM. Org: Online Mendelian Inheritance in Man (OMIM®), an Online Catalog of Human Genes and Genetic Disorders.” Nucleic Acids Research 43 (D1). Oxford Univ Press: D789-98. http://nar.oxfordjournals.org/content/43/D1/D789.short.

Asaoka Yoshinari. 2012. “Phosphorylation of Gli by cAMP-Dependent Protein Kinase.” Vitamins and Hormones 88: 293-307. doi:10.1016/B978-0-12-394622-5.00013-4.

Bachmann Verena A., Anna Riml, Roland G. Huber, George S. Baillie, Klaus R. Liedl, Taras Valovka, and Eduard Stefan. 2013. “Reciprocal Regulation of PKA and Rac Signaling.” Proceedings of the National Academy of Sciences of the United States of America 110 (21): 8531-36. doi:10.1073/pnas.1215902110.

Berger Alice H., Angela N. Brooks, Xiaoyun Wu, Yashaswi Shrestha, Candace Chouinard, Federica Piccioni, Mukta Bagul, et al. 2016. “High-Throughput Phenotyping of Lung Cancer Somatic Mutations.” Cancer Cell 30 (2): 214-28. doi:10.1016/j.ccell.2016.06.022.

Bougen-Zhukov Nicola, Sheng Yang Loh, Hwee Kuan Lee, and Lit-Hsin Loo. 2016. “Large-Scale Image-Based Screening and Profiling of Cellular Phenotypes.” Cytometry. Part A: The Journal of the International Society for Analytical Cytology, July. doi:10.1002/cyto.a.22909.

Boutros Michael, and Julie Ahringer. 2008. “The Art and Design of Genetic Screens: RNA Interference.” Nature Reviews. Genetics 9 (7): 554-66. doi:10.1038/nrg2364.

Bray Mark-Anthony, Adam N. Fraser, Thomas P. Hasaka, and Anne E. Carpenter. 2012. “Workflow and Metrics for Image Quality Control in Large-Scale High-Content Screens.” Journal of Biomolecular Screening 17 (2): 266-74. doi:10.1177/1087057111420292.

Bray Mark-Anthony, Shantanu Singh, Han Han, Chadwick T. Davis, Blake Borgeson, Cathy Hartland, Maria Kost-Alimova, Sigrun M. Gustafsdottir, Christopher C. Gibson, and Anne E. Carpenter. 2016. “Cell Painting, a High-Content Image-Based Assay for Morphological Profiling Using Multiplexed Fluorescent Dyes.” Nature Protocols 11 (9): 1757-74. doi:10.1038/nprot.2016.105.

Caicedo Juan C., Shantanu Singh, and Anne E. Carpenter. 2016. “Applications in Image-Based Profiling of Perturbations.” Current Opinion in Biotechnology 39 (June): 134-42. doi:10.1016/j.copbio.2016.04.003.

Cantwell-Dorris Emma R., John J. O’Leary, and Orla M. Sheils. 2011. “BRAFV600E: Implications for Carcinogenesis and Molecular Therapy.” Molecular Cancer Therapeutics 10 (3): 385-94. doi:10.1158/1535-7163.MCT-10-0799.

Carpenter Anne E., Thouis R. Jones, Michael R. Lamprecht, Colin Clarke, In Han Kang, Ola Friman, David A. Guertin, et al. 2006. “CellProfiler: Image Analysis Software for Identifying and Quantifying Cell Phenotypes.” Genome Biology 7 (10): R100. doi:10.1186/gb2006-7-10-r100.

Cheung Lydia Wt, and Gordon B. Mills. 2016. “Targeting Therapeutic Liabilities Engendered by PIK3R1 Mutations for Cancer Treatment.” Pharmacogenomics 17 (3): 297-307. doi:10.2217/pgs.15.174.

Chircop Megan. 2014. “Rho GTPases as Regulators of Mitosis and Cytokinesis in Mammalian Cells.” Small GTPases 5 (July). doi:10.4161/sgtp.29770.

Cho Hyun Hwa, Keun Koo Shin, Yeon Jeong Kim, Ji Sun Song, Jong Myung Kim, Yong Chan Bae, Chi Dae Kim, and Jin Sup Jung. 2010. “NF-$\kappa$B Activation Stimulates Osteogenic Differentiation of Mesenchymal Stem Cells Derived from Human Adipose Tissue by Increasing TAZ Expression.” Journal of Cellular Physiology 223 (1). Wiley Online Library: 168-77. http://onlinelibrary.wiley.com/doi/10.1002/jcp.22024/full.

Chung Namjin, Xiaohua Douglas Zhang, Anthony Kreamer, Louis Locco, Pei-Fen Kuan, Steven Bartz, Peter S. Linsley, Marc Ferrer, and Berta Strulovici. 2008. “Median Absolute Deviation to Improve Hit Selection for Genome-Scale RNAi Screens.” Journal of Biomolecular Screening 13 (2): 149-58. doi:10.1177/1087057107312035.

Davies Helen, Graham R. Bignell, Charles Cox, Philip Stephens, Sarah Edkins, Sheila Clegg, Jon Teague, et al. 2002. “Mutations of the BRAF Gene in Human Cancer.” Nature 417 (6892): 949-54. doi:10.1038/nature00766.

Davies M. A., K. Stemke-Hale, C. Tellez, T. L. Calderone, W. Deng, V. G. Prieto, A. J. F. Lazar, J. E. Gershenwald, and G. B. Mills. 2008. “A Novel AKT3 Mutation in Melanoma Tumours and Cell Lines.” British Journal of Cancer 99 (8): 1265-68. doi:10.1038/sj.bjc.6604637.

Duda R. O., P. E. Hart, and D. G. Stork. 2012. Pattern Classification. Wiley. https://books.google.com/books?id=Br33IRC3PkQC.

Dupont Sirio, Leonardo Morsut, Mariaceleste Aragona, Elena Enzo, Stefano Giulitti, Michelangelo Cordenonsi, Francesca Zanconato, et al. 2011. “Role of YAP/TAZ in Mechanotransduction.” Nature 474 (7350): 179-83. doi:10.1038/nature10137.

Elsum Imogen A., Claire Martin, and Patrick O. Humbert. 2013. “Scribble Regulates an EMT Polarity Pathway through Modulation of MAPK-ERK Signaling to Mediate Junction Formation.” Journal of Cell Science 126 (Pt 17): 3990-99. doi:10.1242/jcs.129387.

Eser, S., A. Schnieke, G. Schneider, and D. Saur. 2014. “Oncogenic KRAS Signalling in Pancreatic Cancer.” British Journal of Cancer 111 (5): 817-22. doi:10.1038/bjc.2014.215.

Fischer Bernd, Thomas Sandmann, Thomas Horn, Maximilian Billmann, Varun Chaudhary, Wolfgang Huber, and Michael Boutros. 2015. “A Map of Directional Genetic Interactions in a Metazoan Cell.” eLife 4 (March). doi:10.7554/eLife.05464.

Fuchs Florian, Gregoire Pau, Dominique Kranz, Oleg Sklyar, Christoph Budjan, Sandra Steinbrink, Thomas Horn, Angelika Pedal, Wolfgang Huber, and Michael Boutros. 2010. “Clustering Phenotype Populations by Genome-Wide RNAi and Multiparametric Imaging.” Molecular Systems Biology 6 (June): 370. doi:0.1038/msb.2010.25.

Godde Nathan J., Julie M. Sheridan, Lorey K. Smith, Helen B. Pearson, Kara L. Britt, Ryan C. Galea, Laura L. Yates, Jane E. Visvader, and Patrick O. Humbert. 2014. “Scribble Modulates the MAPK/Fra1 Pathway to Disrupt Luminal and Ductal Integrity and Suppress Tumour Formation in the Mammary Gland.” PLoS Genetics 10 (5): e1004323. doi:10.1371/journal.pgen.1004323.

Grech Adrian P., Michelle Amesbury, Tyani Chan, Sandra Gardam, Antony Basten, and Robert Brink. 2004. “TRAF2 Differentially Regulates the Canonical and Noncanonical Pathways of NF-kappaB Activation in Mature B Cells.” Immunity 21 (5): 629-42. doi:10.1016/j.immuni.2004.09.011.

Gustafsdottir S. M., V. Ljosa, K. L. Sokolnicki, J. Anthony Wilson, D. Walpita, M. M. Kemp, K. Petri Seiler, et al. 2013. “Multiplex Cytological Profiling Assay to Measure Diverse Cellular States.” PloS One 8: e80999. doi:10.1371/journal.pone.0080999

Hemmings Brian A., and David F. Restuccia. 2015. “The PI3K-PKB/Akt Pathway.” Cold Spring Harbor Perspectives in Biology 7 (4). doi:10.1101/cshperspect.a026609.

Higuchi, M., N. Masuyama, Y. Fukui, A. Suzuki, and Y. Gotoh. 2001. “Akt Mediates Rac/Cdc42-Regulated Cell Motility in Growth Factor-Stimulated Cells and in Invasive PTEN Knockout Cells.” Current Biology: CB 11 (24):1958-62. https://www.ncbi.nlm.nih.gov/pubmed/11747822.

Hoesel Bastian, and Johannes A. Schmid. 2013. “The Complexity of NF-**κ**B Signaling in Inflammation and Cancer.” Molecular Cancer 12 (August): 86. doi:10.1186/1476-4598-12-86.

Jalili Ahmad, Christine Wagner, Mikhail Pashenkov, Gaurav Pathria, Kirsten D. Mertz, Hans R. Widlund, Mathieu Lupien, et al. 2012. “Dual Suppression of the Cyclin-Dependent Kinase Inhibitors CDKN2C and CDKN1A in Human Melanoma.” Journal of the National Cancer Institute 104 (21): 1673-79. doi:10.1093/jnci/djs373.

Jin Jin, Yichuan Xiao, Hongbo Hu, Qiang Zou, Yanchuan Li, Yanpan Gao, Wei Ge, Xuhong Cheng, and Shao-Cong Sun. 2015. “Proinflammatory TLR Signalling Is Regulated by a TRAF2-Dependent Proteolysis Mechanism in Macrophages.” Nature Communications 6 (January): 5930. doi:10.1038/ncomms6930.

Johnson Randy, and Georg Halder. 2014. “The Two Faces of Hippo: Targeting the Hippo Pathway for Regenerative Medicine and Cancer Treatment.” Nature Reviews. Drug Discovery 13 (1): 63-79. doi:10.1038/nrd4161.

Kandoth Cyriac, Michael D. McLellan, Fabio Vandin, Kai Ye, Beifang Niu, Charles Lu, Mingchao Xie, et al. 2013. “Mutational Landscape and Significance across 12 Major Cancer Types.” Nature 502 (7471): 333-39. doi:10.1038/nature12634.

Kim Eejung, Nina Ilic, Yashaswi Shrestha, Lihua Zou, Atanas Kamburov, Cong Zhu, Xiaoping Yang, et al. 2016. “Systematic Functional Interrogation of Rare Cancer Variants Identifies Oncogenic Alleles.” Cancer Discovery 6 (7): 714-26. doi:10.1158/2159-8290.CD-16-0160.

Kim M. S., E. G. Jeong, N. J. Yoo, and S. H. Lee. 2008. “Mutational Analysis of Oncogenic AKT E17K Mutation in Common Solid Cancers and Acute Leukaemias.” British Journal of Cancer 98 (9): 1533-35. doi:10.1038/sj.bjc.6604212.

Kovacina Kristina S., Grace Y. Park, Sun Sik Bae, Andrew W. Guzzetta, Erik Schaefer, Morris J. Birnbaum, and Richard A. Roth. 2003. “Identification of a Proline-Rich Akt Substrate as a 14-3-3 Binding Partner.” The Journal of Biological Chemistry 278 (12): 10189-94. doi:10.1074/jbc.M210837200.

Lamb Justin, Emily D. Crawford, David Peck, Joshua W. Modell, Irene C. Blat, Matthew J. Wrobel, Jim Lerner, et al. 2006. “The Connectivity Map: Using Gene-Expression Signatures to Connect Small Molecules, Genes, and Disease.” Science 313 (5795): 1929-35. doi:10.1126/science.1132939.

Lang, P., F. Gesbert, M. Delespine-Carmagnat, R. Stancou, M. Pouchelet, and J. Bertoglio. 1996. “Protein Kinase A Phosphorylation of RhoA Mediates the Morphological and Functional Effects of Cyclic AMP in Cytotoxic Lymphocytes.” The EMBO Journal 15 (3):510-19. http://www.ncbi.nlm.nih.gov/pubmed/8599934.

Lee Jongwon, Bang Ung Youn, Kabsun Kim, Jung Ha Kim, Da-Hye Lee, Semun Seong, Inyoung Kim, et al. 2015. “Mst2 Controls Bone Homeostasis by Regulating Osteoclast and Osteoblast Differentiation.” Journal of Bone and Mineral Research: The Official Journal of the American Society for Bone and Mineral Research 30 (9): 1597-1607. doi:10.1002/jbmr.2503.

Leonetti Manuel D., Sayaka Sekine, Daichi Kamiyama, Jonathan S. Weissman, and Bo Huang. 2016. “A Scalable Strategy for High-Throughput GFP Tagging of Endogenous Human Proteins.” Proceedings of the National Academy of Sciences of the United States of America 113 (25): E3501-8. doi:10.1073/pnas.1606731113.

Liu Bo, Yonggang Zheng, Feng Yin, Jianzhong Yu, Neal Silverman, and Duojia Pan. 2016. “Toll Receptor-Mediated Hippo Signaling Controls Innate Immunity in Drosophila.” Cell 164 (3): 406-19. doi:10.1016/j.cell.2015.12.029.

Liu Chen-Ying, Xianbo Lv, Tingting Li, Yanping Xu, Xin Zhou, Shimin Zhao, Yue Xiong, Qun-Ying Lei, and Kun-Liang Guan. 2011. “PP1 Cooperates with ASPP2 to Dephosphorylate and Activate TAZ.” The Journal of Biological Chemistry 286 (7): 5558-66. doi:10.1074/jbc.M110.194019.

Ljosa Vebjorn, Peter D. Caie, Rob Ter Horst, Katherine L. Sokolnicki, Emma L. Jenkins, Sandeep Daya, Mark E. Roberts, et al. 2013. “Comparison of Methods for Image-Based Profiling of Cellular Morphological Responses to Small-Molecule Treatment.” Journal of Biomolecular Screening 18 (10): 1321-29. doi:10.1177/1087057113503553.

Maddika Subbareddy, Sudharsana Rao Ande, Emilia Wiechec, Lise Lotte Hansen, Sebastian Wesselborg, and Marek Los. 2008. “Akt-Mediated Phosphorylation of CDK2 Regulates Its Dual Role in Cell Cycle Progression and Apoptosis.” Journal of Cell Science 121 (Pt 7): 979-88. doi:10.1242/jcs.009530.

Marivin, A., J. Berthelet, J. Cartier, C. Paul, S. Gemble, A. Morizot, W. Boireau, et al. 2014. “cIAP1 Regulates TNF-Mediated cdc42 Activation and Filopodia Formation.” Oncogene 33 (48): 5534-45. doi:10.1038/onc.2013.499.

Martin Sophie G. 2015. “Spontaneous Cell Polarization: Feedback Control of Cdc42 GTPase Breaks Cellular Symmetry.” BioEssays: News and Reviews in Molecular, Cellular and Developmental Biology 37 (11): 1193-1201. doi:10.1002/bies.201500077.

McCOY Melissa S., Cornelia I. Bargmann, and Robert A. Weinberg. 1984. “Human Colon Carcinoma Ki-ras2 Oncogene and Its Corresponding Proto-Oncogene.” Molecular and Cellular Biology 4 (8). Am Soc Microbiol: 1577-82. http://mcb.asm.org/content/4/8/1577.short.

Melendez Jaime, Matthew Grogg, and Yi Zheng. 2011. “Signaling Role of Cdc42 in Regulating Mammalian Physiology.” The Journal of Biological Chemistry 286 (4): 2375-81. doi:10.1074/jbc.R110.200329.

Meng Zhipeng, Toshiro Moroishi, and Kun-Liang Guan. 2016. “Mechanisms of Hippo Pathway Regulation.” Genes & Development 30 (1): 1-17. doi:10.1101/gad.274027.115.

Mohseni Morvarid, Jianlong Sun, Allison Lau, Stephen Curtis, Jeffrey Goldsmith, Victor L. Fox, Chongjuan Wei, et al. 2014. “A Genetic Screen Identifies an LKB1-MARK Signalling Axis Controlling the Hippo-YAP Pathway.” Nature Cell Biology 16 (1): 108-17. doi:10.1038/ncb2884.

Mukherji Mridul, Russell Bell, Lubica Supekova, Yan Wang, Anthony P. Orth, Serge Batalov, Loren Miraglia, et al. 2006. “Genome-Wide Functional Analysis of Human Cell-Cycle Regulators.” Proceedings of the National Academy of Sciences of the United States of America 103 (40): 14819-24. doi:10.1073/pnas.0604320103.

Murai Kasumi, and Richard Treisman. 2002. “Interaction of Serum Response Factor (SRF) with the Elk-1 B Box Inhibits RhoA-Actin Signaling to SRF and Potentiates Transcriptional Activation by Elk-1.” Molecular and Cellular Biology 22 (20):7083-92. https://www.ncbi.nlm.nih.gov/pubmed/12242287.

Nobes C. D., and A. Hall. 1999. “Rho GTPases Control Polarity, Protrusion, and Adhesion during Cell Movement.” The Journal of Cell Biology 144 (6):1235-44. https://www.ncbi.nlm.nih.gov/pubmed/10087266.

Oishi Atsuro, Noriko Makita, Junichiro Sato, and Taroh Iiri. 2012. “Regulation of RhoA Signaling by the cAMP-Dependent Phosphorylation of RhoGDIα.” The Journal of Biological Chemistry 287 (46): 38705-15. doi:10.1074/jbc.M112.401547.

Pau Gregoire, Thomas Walter, Beate Neumann, Jean-Karim Hériché, Jan Ellenberg, and Wolfgang Huber. 2013. “Dynamical Modelling of Phenotypes in a Genome-Wide RNAi Live-Cell Imaging Assay.” BMC Bioinformatics 14 (October): 308. doi:10.1186/1471-2105-14-308.

Reginensi Antoine, Rizaldy P. Scott, Alex Gregorieff, Mazdak Bagherie-Lachidan, Chaeuk Chung, Dae-Sik Lim, Tony Pawson, Jeff Wrana, and Helen McNeill. 2013. “Yap- and Cdc42-Dependent Nephrogenesis and Morphogenesis during Mouse Kidney Development.” PLoS Genetics 9 (3): e1003380. doi:10.1371/journal.pgen.1003380.

Rolli-Derkinderen Malvyne, Vincent Sauzeau, Laurent Boyer, Emmanuel Lemichez, Céline Baron, Daniel Henrion, Gervaise Loirand, and Pierre Pacaud. 2005. “Phosphorylation of Serine 188 Protects RhoA from Ubiquitin/proteasome-Mediated Degradation in Vascular Smooth Muscle Cells.” Circulation Research 96 (11): 1152-60. doi:10.1161/01.RES.0000170084.88780.ea.

Samaj Jozef, Frantisek Baluska, and Heribert Hirt. 2004. “From Signal to Cell Polarity: Mitogen-activated Protein Kinases as Sensors and Effectors of Cytoskeleton Dynamicity.” Journal of Experimental Botany 55 (395): 189-98. doi:10.1093/jxb/erh012.

Schmich Fabian, Ewa Szczurek, Saskia Kreibich, Sabrina Dilling, Daniel Andritschke, Alain Casanova, Shyan Huey Low, et al. 2015. “gespeR: A Statistical Model for Deconvoluting off-Target-Confounded RNA Interference Screens.” Genome Biology 16 (October): 220. doi:10.1186/s13059-015-0783-1.

Shehu Amarda, Daniel Barbará, and Kevin Molloy. 2016. “A Survey of Computational Methods for Protein Function Prediction.” In Big Data Analytics in Genomics, edited by Ka-Chun Wong, 225-98. Springer International Publishing. doi:10.1007/978-3-319-41279-5_7.

Shin Ilchung, Seonhoe Kim, Hyun Song, Hyeong-Reh Choi Kim, and Aree Moon. 2005. “H-Ras-Specific Activation of Rac-MKK3/6-p38 Pathway: Its Critical Role in Invasion and Migration of Breast Epithelial Cells.” The Journal of Biological Chemistry 280 (15): 14675-83. doi:10.1074/jbc.M411625200.

Singh Shantanu, Xiaoyun Wu, Vebjorn Ljosa, Mark-Anthony Bray, Federica Piccioni, David E. Root, John G. Doench, Jesse S. Boehm, and Anne E. Carpenter. 2015. “Morphological Profiles of RNAi-Induced Gene Knockdown Are Highly Reproducible but Dominated by Seed Effects.” PloS One 10 (7): e0131370. doi:10.1371/journal.pone.0131370.

Stark Chris, Bobby-Joe Breitkreutz, Teresa Reguly, Lorrie Boucher, Ashton Breitkreutz, and Mike Tyers. 2006. “BioGRID: A General Repository for Interaction Datasets.” Nucleic Acids Research 34 (Database issue): D535-39. doi:10.1093/nar/gkj109.

Stengel Kristy R., and Yi Zheng. 2012. “Essential Role of Cdc42 in Ras-Induced Transformation Revealed by Gene Targeting.” PloS One 7 (6): e37317. doi:10.1371/journal.pone.0037317.

Subramanian Aravind, Pablo Tamayo, Vamsi K. Mootha, Sayan Mukherjee, Benjamin L. Ebert, Michael A. Gillette, Amanda Paulovich, et al. 2005. “Gene Set Enrichment Analysis: A Knowledge-Based Approach for Interpreting Genome-Wide Expression Profiles.” Proceedings of the National Academy of Sciences of the United States of America 102 (43): 15545-50. doi:10.1073/pnas.0506580102.

Tada, K., T. Okazaki, S. Sakon, T. Kobarai, K. Kurosawa, S. Yamaoka, H. Hashimoto, et al. 2001. “Critical Roles of TRAF2 and TRAF5 in Tumor Necrosis Factor-Induced NF-Kappa B Activation and Protection from Cell Death.” The Journal of Biological Chemistry 276 (39): 36530-34. doi:10.1074/jbc.M104837200.

Tornatore Laura, Anil K. Thotakura, Jason Bennett, Marta Moretti, and Guido Franzoso. 2012. “The Nuclear Factor Kappa B Signaling Pathway: Integrating Metabolism with Inflammation.” Trends in Cell Biology 22 (11): 557-66. doi:10.1016/j.tcb.2012.08.001.

Varelas Xaralabos. 2014. “The Hippo Pathway Effectors TAZ and YAP in Development, Homeostasis and Disease.” Development 141 (8): 1614-26. doi:10.1242/dev.102376.

Veitia Reiner A. 2007. “Exploring the Molecular Etiology of Dominant-Negative Mutations.” The Plant Cell 19 (12): 3843-51. doi:10.1105/tpc.107.055053.

Wawer Mathias J., Kejie Li, Sigrun M. Gustafsdottir, Vebjorn Ljosa, Nicole E. Bodycombe, Melissa A. Marton, Katherine L. Sokolnicki, et al. 2014. “Toward Performance-Diverse Small-Molecule Libraries for Cell-Based Phenotypic Screening Using Multiplexed High-Dimensional Profiling.” Proceedings of the National Academy of Sciences of the United States of America 111 (30): 10911-16. doi:10.1073/pnas.1410933111.

Wiza Claudia, Alexandra Chadt, Marcel Blumensatt, Timo Kanzleiter, Daniella Herzfeld De Wiza, Angelika Horrighs, Heidi Mueller, et al. 2014. “Over-Expression of PRAS40 Enhances Insulin Sensitivity in Skeletal Muscle.” Archives of Physiology and Biochemistry 120 (2): 64-72. doi:10.3109/13813455.2014.894076.

Wu Xue, Jeremy Simpson, Jenny H. Hong, Kyoung-Han Kim, Nirusha K. Thavarajah, Peter H. Backx, Benjamin G. Neel, and Toshiyuki Araki. 2011. “MEK-ERK Pathway Modulation Ameliorates Disease Phenotypes in a Mouse Model of Noonan Syndrome Associated with the Raf1L613V Mutation.” doi:10.1172/JCI44929.

Yang Xiaoping, Jesse S. Boehm, Xinping Yang, Kourosh Salehi-Ashtiani, Tong Hao, Yun Shen, Rakela Lubonja, et al. 2011. “A Public Genome-Scale Lentiviral Expression Library of Human ORFs.” Nature Methods 8 (8): 659-61. doi:10.1038/nmeth.1638.

Yu Guangchuang, Li-Gen Wang, Yanyan Han, and Qing-Yu He. 2012. “clusterProfiler: An R Package for Comparing Biological Themes among Gene Clusters.” Omics: A Journal of Integrative Biology 16 (5): 284-87. doi:10.1089/omi.2011.0118.

Zhang, S., J. Han, M. A. Sells, J. Chernoff, U. G. Knaus, R. J. Ulevitch, and G. M. Bokoch. 1995. “Rho Family GTPases Regulate p38 Mitogen-Activated Protein Kinase through the Downstream Mediator Pak1.” The Journal of Biological Chemistry 270 (41):23934-36. https://www.ncbi.nlm.nih.gov/pubmed/7592586.

Zhang Z-G, C. A. Lambert, S. Servotte, G. Chometon, B. Eckes, T. Krieg, C. M. Lapière, B. V. Nusgens, and M. Aumailley. 2006. “Effects of Constitutively Active GTPases on Fibroblast Behavior.” Cellular and Molecular Life Sciences: CMLS 63 (1): 82-91. doi:10.1007/s00018-005-5416-5.

Zhao Bin, Xin Ye, Jindan Yu, Li Li, Weiquan Li, Siming Li, Jianjun Yu, et al. 2008. “TEAD Mediates YAP-Dependent Gene Induction and Growth Control.” Genes & Development 22 (14): 1962-71. doi:10.1101/gad.1664408.

